# Enhanced lung delivery of an immunostimulatory duplex RNA augments the antitumor activity by reshaping systemic cytokine pharmacodynamics

**DOI:** 10.64898/2026.05.03.722518

**Authors:** Eliz Amar-Lewis, Alexander M. Cryer, Jie Ji, Chaitra Belgur, Anastasia Ershova, Nelly Andrews Interiano, William Sawyer, Zohar Pode, Namrata Ramani, Juan Carlos Oliva Estrada, Nathalie Nicole Casteele Hernandez, John F.K Sauld, Yuncheng Man, Sylvie G. Bernier, Amanda R. Graveline, Melinda Sanchez Suarez, Girija Goyal, Kenneth E. Carlson, William M. Shih, Donald E. Ingber, Natalie Artzi

**Affiliations:** Wyss Institute for Biologically Inspired Engineering, Harvard University, Boston, MA 02215; Institute for Medical Engineering and Science, Massachusetts Institute of Technology, Cambridge, MA 02139; Department of Medicine, Division of Engineering in Medicine, Brigham and Women’s Hospital, Harvard Medical School, Boston, MA 02115; Department of Cancer Biology, Dana-Farber Cancer Institute, Boston, MA, 02215; Department of Biological Chemistry and Molecular Pharmacology, Harvard Medical School, Boston, MA, 02115; Vascular Biology Program, Boston Children’s Hospital and Harvard Medical School, Boston, MA 02115; Harvard John A. Paulson School of Engineering and Applied Sciences, Harvard University, Boston, MA 02134

## Abstract

The organ-specific enrichment of drug delivery vehicles, such as lipid nanoparticles (LNPs), can be leveraged to concentrate drugs at disease sites to increase efficacy and limit toxicity. For immunostimulatory therapeutics, however, tissue accumulation beyond diseased sites may also shape drug activity by determining which organs and cell populations first sense the agonist and initiate downstream immune responses. Here, we show that the anticancer efficacy of an immunostimulatory duplex RNA (dsRNA) can be augmented using LNPs that are formulated to preferentially target the lung, which dictates the systemic pharmacodynamics of the cytokines it elicits. The immunostimulatory dsRNA was formulated into LNPs engineered for either enhanced liver-(LiverLNPs) or lung-(LungLNPs) based delivery, matched for size, encapsulation efficiency, and in vitro potency.

In mice, delivery of dsRNA in LungLNPs enhanced uptake into endothelial, epithelial, and resident immune cells populations and induced substantially higher circulating levels of type I, type III interferons and proinflammatory cytokines than dsRNA formulated in LiverLNPs. This significant systemic response induced by lung-enhanced delivery required competent retinoic acid-inducible gene I and Toll-like receptor 7 signaling. Functionally, LNPs that preferentially targeted the lungs induced significantly greater suppression of tumor growth in both subcutaneous and metastatic models of melanoma. LungLNP/dsRNA also induced cytokine secretion and inhibited tumor cell proliferation in a human lung cancer-on-a-chip model. Together, these results establish that pulmonary exposure can alter systemic pharmacodynamics and therapeutic activity of immunostimulatory RNA.

## Introduction

The organ-specific enrichment of a drug is often framed as a way to concentrate the therapeutic at the main organ that harbors the disease^1–4^. For immunostimulatory RNA, however, organ tropism may also alter the systemic pharmacodynamics (PD) of the immunomodulatory molecules it elicits, depending on which cell types sense the RNA after systemic dosing. In this context, delivery of an immunostimulatory RNA to specific organ could influence anticancer activity at that site where it accumulates or by altering the magnitude and timing of the circulating cytokine responses it elicits.

The recognition of viral RNAs by the innate immune system is an evolutionarily conserved defense mechanism that initiates the rapid production of interferons (IFNs), cytokines, and chemokines that restrict viral replication and promote antitumor immunity ^5,6^. Such foreign-RNA sensing is mediated by pattern-recognition receptors that detect distinct RNA features and initiate transcriptional programs of self-defense, including the expression of IFN-stimulated genes ^7–9^. Consequently, activation of RNA sensors, such as retinoic acid-inducible gene-I (RIG-I)^10–12^ and Toll-like receptors (TLRs)^13–16^ is being explored as an effective approach in cancer immunotherapy^12,17–20^.

Mucosal tissues in organs that are exposed to the external environment, such as the lung, create a specialized immune environment that serves as a frontline barrier to pathogens by expressing high levels of innate nucleic acid sensors^21–23^, with defined spatial distribution in the context of RNA sensing^24^. Recent work with human cells and a human lung-on-a-chip (Lung Chip) microfluidic culture model led to the identification of short double stranded RNAs (dsRNAs) with potent immunostimulatory activity that can induce type-I and type-III IFNs through RIG-I-dependent signaling^25^. These dsRNAs are non-canonical agonists that do not require 5’ tri-phosphorylation^26–28^ to activate RIG-I and which exert broad spectrum antiviral activity against strains of influenza and coronaviruses in human cells in vitro and afforded significant antiviral activity against viral infection in a mouse SARS-CoV2 model. Given that IFN-α has been approved by the U.S. Food and Drug Administration (FDA) for the treatment of certain cancers^29–32^, and transient IFN production can augment antigen presentation and cytotoxic function in experimental cancer models^33–35^, we set out to examine whether a model immunostimulatory dsRNA (RNA-1) also exerts anti-cancer activity. While the dsRNA was previously shown to be active when administered systemically (intravenously) in mice infected with SARS-CoV-2^36–38^, use of drug delivery vehicles, such as lipid nanoparticles (LNPs), substantially enhances the stability and efficacy of RNA therapies^39–44^. Importantly, by altering the tissue and cell distribution of the immunostimulatory dsRNA using LNPs engineered to modify organ tropism^45–48^, it also may be possible to reshape intrinsic inflammatory potential and intracellular trafficking, thereby shifting the PD of the elicited immunomodulatory responses. Unravelling these effects is necessary to correlate between agonist biodistribution (BD), innate sensing and therapeutic efficacy.

Thus, in this study, we tested the impact of LNP formulation of an immunostimulatory dsRNA on cellular and organ distribution, in vivo PD, and cancer efficacy. We formulated RNA-1 in a clinically relevant LNP formulation that delivers RNA cargo primarily to the liver (LiverLNPs) or a chemically modified LNP formulation with enhanced delivery of RNA cargo to the lung (LungLNPs)^46,49,50^ and compared in vitro activity, in vivo plasma secretion of cytokines, biodistribution, immune phenotyping analysis, and antitumor efficacy. We also tested the innate receptor specificity induced by RNA-1 in the plasma-cytokine response (Figure 1a).

**Fig. 1.**
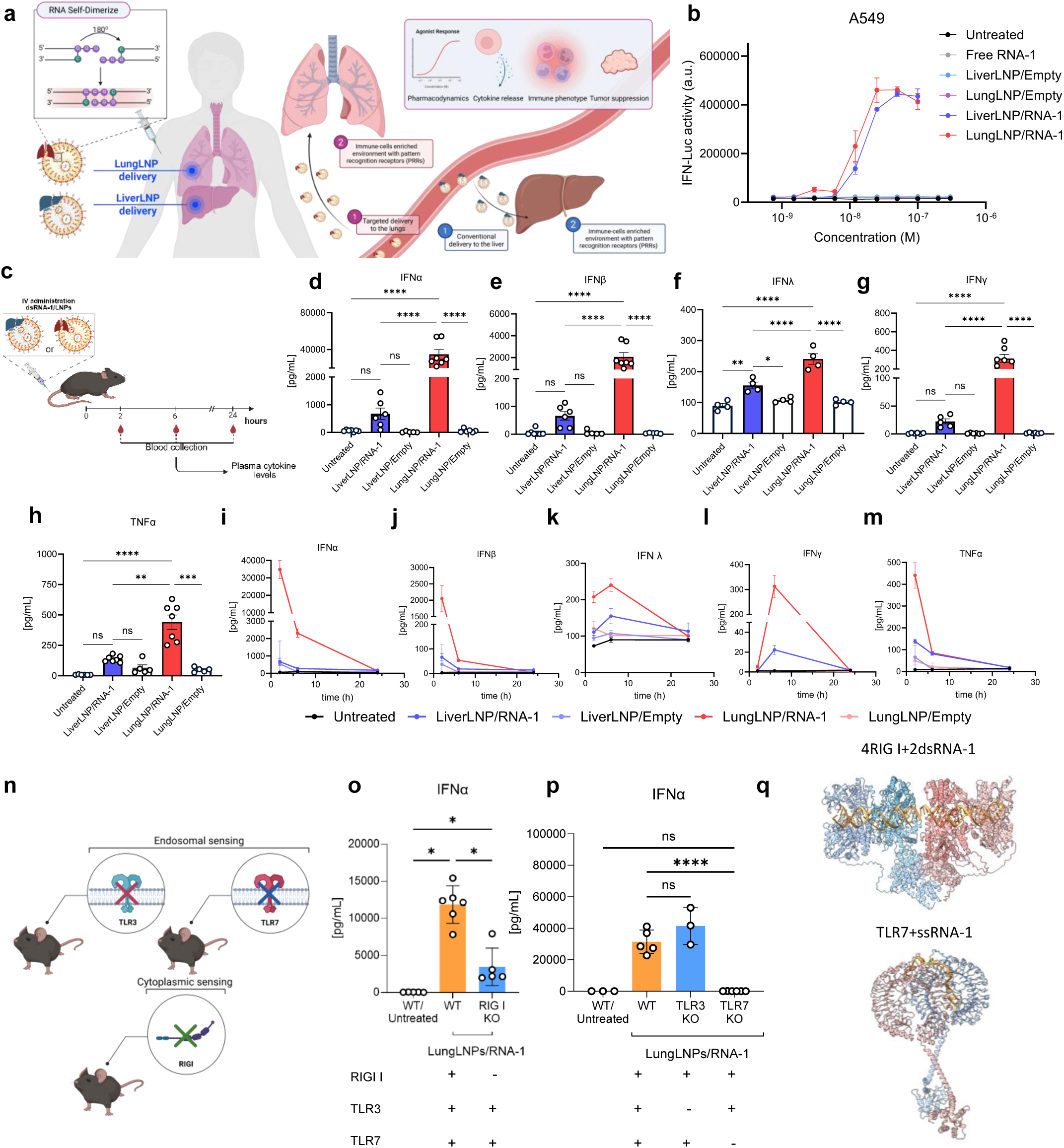
Lung-tropic delivery of RNA-1 reveals organ-specific innate immune activation and systemic cytokine responses dependent on RIG I and TLR7 in mice. (a) Schematic illustrating how BD shapes the in vivo immunostimulatory activity of self-dimerizing RNA-1 delivered by LungLNPs or LiverLNPs. LungLNP enhances delivery of RNA-1 to the lungs (1, pink), whereas conventional LiverLNP delivery directs RNA-1 to the liver (1, blue). In each case, organ-specific accumulation leads to uptake of RNA-1 into tissue resident immune or non-immune cell populations expressing pattern recognition receptors (PRRs) (2, pink/blue), thereby influencing pharmacodynamic responses, cytokine release, immune activation, and tumor suppression. (b) IFN-luciferase reporter assay in A549 IRF3 dual reporter cells showing induction by RNA-1 formulated in LungLNPs vs LiverLNPs, compared with free RNA-1 and empty controls. Data presented as average ± SD, n = 3. (c) Schematic of the *in vivo* pharmacodynamic (PD) model used to assess plasma cytokines following systemic administration of LungLNP/RNA-1, LiverLNP/RNA-1 formulations and corresponding empty LNPs. Mice were dosed with 2.2 mg/kg of RNA-1. (d-h) Quantification of peak plasma cytokine levels (2h for IFNα, IFNβ, TNFα and 6h for IFNγ, IFNλ), (i–m) Temporal kinetics of plasma cytokines (IFNα, IFNβ, IFNλ, IFNγ, and TNFα) following treatment at 2, 6 and 24 h. Data are represented as mean ± SEM from a representative experiment of three independent experiments with n = 6–7 (d–h) and n = 5–7 (i–m) biologically independent samples. (n) Schematic presentation depicting the knockout models used to study the innate immune pathway activated by LungLNPs/RNA-1 (2.2 mg/kg) in mice. (o) Quantification of IFNα plasma levels in RIG I KO mice (cytoplasmic sensing) compared with wildtype (WT) control. (p) Quantification of IFNα plasma levels in TLR3 and TLR7 KO mice compared with wildtype (WT) control. Data are represented as mean ± SD from a representative experiment of two independent experiments with n = 3-6 (o–p) biologically independent samples. (q) Molecular illustration depicting an Alphafold3 modeling of mouse RIG I and mouse TLR7 engaged with dsRNA-1 or ssRNA-1 respectively. Panels a, c and n were created with BioRender.com. The data were analyzed by ordinary one-way ANOVA with Tukey’s multiple-comparisons test; * *P* ≤ 0.05, ** *P* ≤ 0.01, *** *P* ≤ 0.001, **** *P* ≤ 0.0001.

## Results

### LungLNPs delivery of RNA-1 shapes systemic cytokine pharmacodynamics

To compare how BD delivery affects RNA-1 activity, we encapsulated RNA-1 in LiverLNPs (LiverLNP/RNA-1) and LungLNPs (LungLNP/RNA-1), which had similar particle size (110–130 nm), polydispersity (0.10–0.16), and encapsulation efficiency (>93%), yet differed in zeta potential, consistent with the intended design (**Supplementary Fig. 1a–c**)^49,50^. Both formulations induced the activity of interferon regulatory factor 3 (IRF3) in cultured A549 reporter cells with similar EC_50_ values (LungLNP/RNA-1, 12.1 nM; LiverLNP/RNA-1, 15.4 nM) without compromising cell viability in vitro (**Fig. 1b; Supplementary Fig. 2**). In cultures of primary bone marrow-derived dendritic cells, both LiverLNP/RNA-1 and LungLNP/RNA-1 increased the expression of co-stimulatory markers (CD40, CD80, and CD86) and major histocompatibility complex class II (MHC II) compared with empty LNPs and untreated controls, with comparable magnitude between formulations (**Supplementary Fig. 3a–d**). Together, these data demonstrate that the two LNP formulations exhibit similar potency in vitro. In vivo cytokine responses were determined in C57BL/6 mice following intravenous administration of either LungLNP/RNA-1 or LiverLNP/RNA-1 or their respective empty-LNP controls (all at 2.2 mg/kg). Plasma was collected at 2, 6 and 24 hours (h) and interferon and cytokine levels were determined (**Fig. 1c**). LungLNP/RNA-1 produced significantly higher circulating levels of Type I and III interferons IFNα, IFNβ, and IFNλ, as well as the Type II interferon IFN-γ and TNFα, than LiverLNP/RNA-1. (**Fig. 1d–h** Looking across the PD response over time, the plasma levels of IFN *α*, IFN *β*, and TNF *α* peaked early and declined toward baseline by 6 h, consistent with a rapid induction and clearance typical of innate cytokine responses, whereas IFN *γ* and IFN *λ* peaked later at 6 h before declining by 24 h, reflecting its secondary induction by activated lymphocytes (**Fig. 1i–m**)^51,52^. We observed that these effects were independent of the vehicle type, as long as there is pulmonary tropism, as intravenous delivery of RNA-1 with JetPEI, a polymeric carrier that preferentially transfects cells in the lung^53–57^, also led to pronounced plasma cytokine production, and these effects were lost upon intraperitoneal administration of RNA-1/JetPEI, as is the pulmonary tropism of this delivery vehicle (**Supplementary Fig. 4**). These results are consistent with pulmonary delivery shaping the PD of the systemic cytokine secretion in response to RNA-1.

### RNA-1–induced cytokine responses are dependent on RIG-I and TLR7

Given that RNA-1 is a non-modified small dsRNA that is recognized by PRRs, we next examined the innate pathways required for the LungLNP-induced interferon response in plasma of mice by administering LungLNP/RNA-1 to mice deficient in key receptors that mediate RNA sensing in the cytosol (RIG-I) and endosome (TLR3 and TLR7) (**Fig. 1n**). In RIG-I knockout mice, LungLNP/RNA-1 produced lower plasma interferon and cytokine levels than in controls (**Fig. 1o; Supplementary Fig. 5a–c**). While the interferon and cytokine response was preserved in TLR3 knockout mice, induction was strongly reduced to basal levels in TLR7-deficient mice (**Fig. 1p; Supplementary Fig. 5d–f**).

Alphafold modeling (**Fig. 1q; Supplementary Fig. 6**) of RNA-1 structure is consistent with a structural arrangement in which two RNA-1 molecules engage with four RIG-I units, potentially stabilized by G-quadruplex-mediated dimerization as previously described^25^. TLR7 is known to recognize ssRNA structures enriched in GU motifs, suggesting that RNA-1 may may be present as ssRNA species inside endosomes, which are sufficient to activate TLR7 as demonstrated with unmodified siRNAs^58,59^ and discussed in **Supplementary note 1**. The complete abrogation of RNA-1-induced cytokine responses in TLR7 knockout mice, with only partial reduction in RIG-I knockout, suggests a dominant role for TLR7 when RNA-1 is delivered systemically using LungLNPs, which is discussed in detail in **Supplementary note 2**. Overall, these data support a model in which RNA-1 engages both RIG-I and TLR7 in mice, with TLR7 as a dominant requirement for the observed plasma cytokine response. These results expand on prior findings in human lung epithelial cells, where RNA-1 induced RIG-I-dependent IRF3 activation, and indicate that murine responses to RNA-1 involves both RIG-I and TLR7 pathways when delivered in LungLNPs.

### RNA-1-enriched lungs induce lung-specific immune activation

To correlate delivery to organ and cellular BD, we performed RNA-1 BD and early PD profiling in B16-F10 tumor-bearing mice using Cy5-labeled RNA-1 (**Fig. 2a**). IVIS live animal imaging at 4 h showed that LiverLNP/RNA-1 primarily delivered RNA-1 to the liver and spleen^60^, whereas LungLNP/RNA-1 preferentially redirected its delivery to the lung while retaining substantial liver and spleen Cy5 RNA-1 fluorescent intensity (**Fig. 2b–d**).

**Fig. 2.**
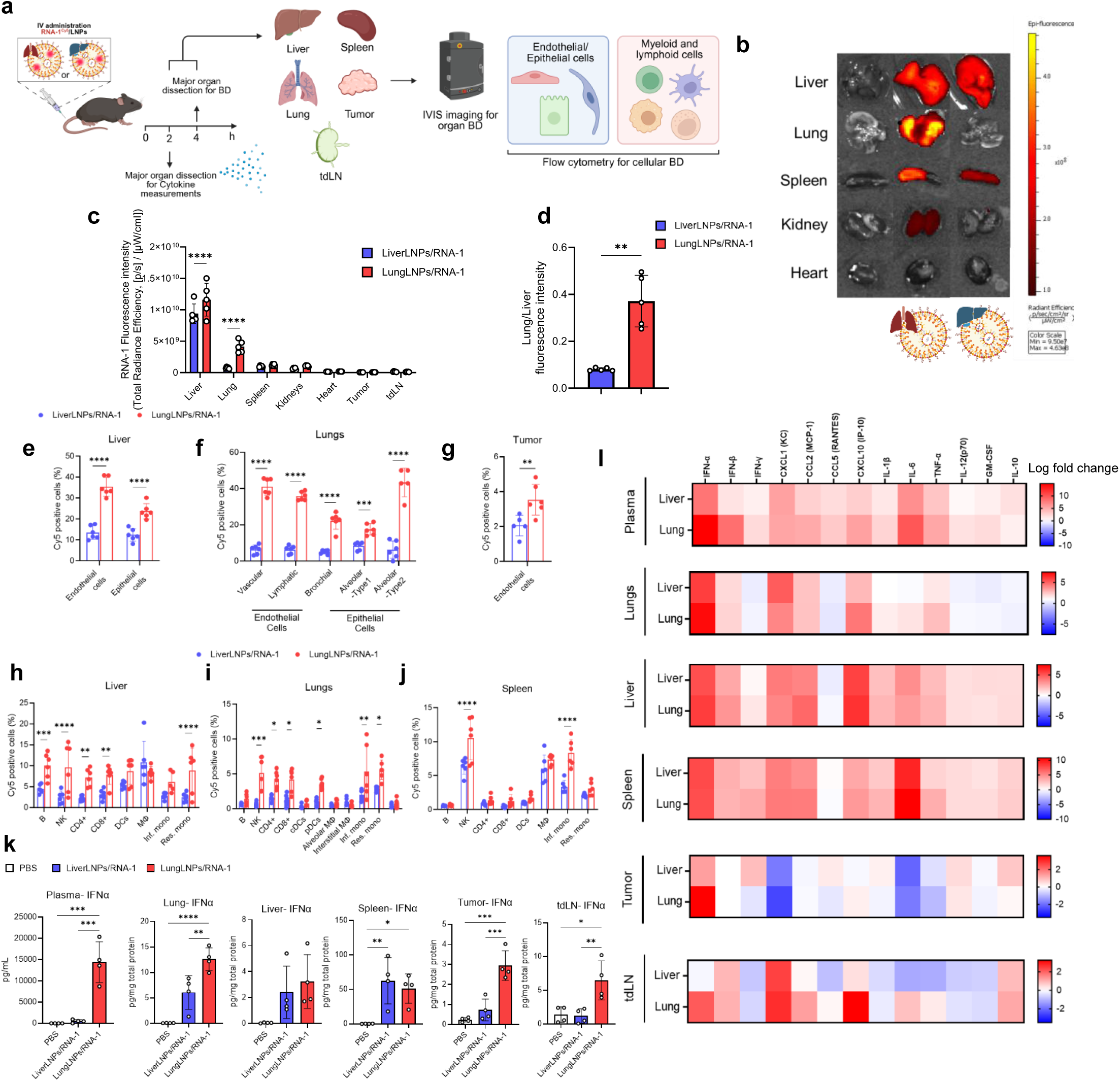
Altering LNP formulation of RNA-1 changes BD, cellular uptake, and immune activation. (a) Schematic of the experimental design for BD and immune activation studies following i.v. administration of Cy5 labeled LungLNP/RNA-1 or LiverLNP/RNA-1 (2.2 mg/kg RNA-1) in mice bearing B16-F10 tumors. Organs were collected 2 h post-injection for cytokine analysis and 4 h post-injection for BD at organ and cellular levels (immune and endothelial/epithelial). (b) Representative IVIS images of organs. (c) Quantification of RNA-1^Cy5^ distribution across organs by IVIS imaging. Data are represented as mean ± SD from a representative experiment of two independent experiments with n = 5, data were analyzed by two-way ANOVA with Tukey’s multiple-comparisons test. (d) Lung:liver organ fluorescence intensity ratio across treatments. Data are represented as mean ± SD with n = 5, data were analyzed by two-tailed Welch’s t test. (e-j) Flow cytometry analysis of cellular internalization in different tissues and cell populations: endothelial and epithelial cells in liver (e), lung (f), and tumor (g), immune cells in liver (h), lung (i), and spleen (j). Heatmap of cytokine levels in lung homogenates 2 h post-treatment, showing strong induction of type I interferons and proinflammatory cytokines in LungLNP/RNA-1–treated animals compared with PBS or LiverLNP/RNA-1. Data are shown for IFNα (k) and full inflammation cytokine panel (l) in the plasma, lung, liver, spleen, tumor and tdLN. Data are presented as mean ± SD with n = 4 biologically independent samples per group. Statistical analyses were performed using one-way ANOVA with Tukey’s multiple-comparisons test. Panel a was created with BioRender.com. * *P* ≤ 0.05, ** *P* ≤ 0.01, *** *P* ≤ 0.001, **** *P* ≤ 0.0001.

Cellular uptake of RNA-1 was determined by flow cytometry in liver, lung, spleen, tumor and tumor-draining lymph node (tdLN) tissues dissociated from animals injected intravenously with RNA-1/LNPs, quantifying the percentage of Cy5-positive cells across immune, endothelial and epithelial populations (**Fig. 2a**). In endothelial and epithelial cells, LiverLNPs showed low RNA-1 uptake (≤10% Cy5-positive) across liver, lung and tumor tissues (**Fig. 2e–g**). In contrast, LungLNP/RNA-1 increased uptake in endothelial and epithelial populations in the liver (35% and 25% Cy5-positive endothelial and epithelial cells, respectively) and lung, including vascular and lymphatic endothelial cells (35–40% Cy5-positive), bronchial epithelial cells (25% Cy5-positive) and alveolar epithelial populations (**Fig. 2e,f**). Type II alveolar epithelial cells, which express innate immune receptors and actively produce type I and III IFNs in response to viral RNA, showed 40% Cy5-positive cells. In tumors grown subcutaneously, LungLNP/RNA-1 increased endothelial uptake relative to LiverLNPs, although the magnitude was smaller than in major organs (**Fig. 2g**).

Immune cell uptake also differed by formulation. In the liver, LiverLNP-enabled uptake of RNA-1 was most apparent in macrophages (10% Cy5-positive) and dendritic cells (5% Cy5-positive), with other immune populations showing ≤5% Cy5-positive cells (**Fig. 2h**). The significant Cy5 fluorescent signal observed in Liver with LiverLNP in whole animal imaging (**Fig. 2b-d**) is probably associated with efficient uptake into hepatocytes, which is known for this formulation ^61–63^. In contrast, LungLNP/RNA-1 increased RNA-1 uptake across additional myeloid and lymphoid cells, including inflammatory monocytes (5% Cy5-positive), resident monocytes (10% Cy5-positive), NK cells (10% Cy5-positive), B cells (10% Cy5-positive), and CD4⁺ and CD8⁺ T cells (7.5% Cy5-positive). In the lungs, LungLNP//RNA-1 yielded detectable uptake across several immune populations, including NK cells, inflammatory and resident monocytes, plasmacytoid dendritic cells, and CD4⁺ and CD8⁺ T cells, (5% Cy5-positive) whereas LiverLNPs showed minimal uptake (**Fig. 2i**). In the spleen, uptake enabled by LungLNP/RNA-1 was most apparent in NK cells (10% Cy5-positive), macrophages (7.5% Cy5-positive) and inflammatory monocytes (7.5% Cy5-positive) (**Fig. 2j**). We did not detect substantial uptake in immune cells from tumor or tdLN under these conditions (**Supplementary Fig. 7**).

To further elucidate the BD-activity relationship of RNA-1, we measured interferons and cytokines in plasma and tissue homogenates 2 h after systemic dosing (**Fig. 2k–l**). Compared with LiverLNP/RNA-1, LungLNP/RNA-1 increased IFN *α* levels 27-fold in plasma and in lung, tumor, and tdLN tissues (2-, 4-, and 5.5-fold, respectively), whereas liver and spleen homogenates showed similar IFN *α* levels between formulations at this time point (**Fig. 2k**). Across a broader panel of analytes, LungLNP/RNA-1 increased interferons, cytokines and chemokines across multiple organs, with prominent responses in lung homogenates (**Fig. 2l**). In particular, high levels of IFN *β*, TNF *α*, interleukin 6 (IL-6) and CXCL10 were induced in the lung, consistent with a proinflammatory environment. LiverLNP/RNA-1 produced localized cytokine responses of lower magnitude and type I and III IFNs remained near basal levels. Increased levels of cytokines and chemokines were observed in the tumor microenvironment and tdLN tissue with LungLNP/RNA-1, including higher levels of proinflammatory cytokines (TNF *α*, IL-6) and chemokines (CXCL-1, CCL2 and CXCL-10)-cytokines associated with immune-cell infiltration and T-cell activation.

### LungLNPs/RNA-1 delivery leads to antitumor activity in primary and metastatic B16-F10 models

It was important to determine if the significant difference in PD response between LungLNP/RNA-1 and LiverLNP/RNA-1 translated to differences in antitumor activity. C57BL/6 mice bearing subcutaneous B16-F10 tumors received three systemic doses of RNA-1 (2.2 mg/kg) formulated in LungLNPs or LiverLNPs, administered once every four days, alone or in combination with anti-PD-1 (100µg/mouse) compared to controls of PBS, empty-LNPs, and anti-PD-1 alone (**Fig. 3a**). LungLNP/RNA-1 significantly slowed tumor growth compared to controls, restricting tumor expansion up to 14 days after the first dose, whereas LiverLNP/RNA-1 had no therapeutic effect in this model showing tumor growth comparable to PBS and empty LNP-treated controls (**Fig. 3b,d–i**). Combination of anti-PD-1 with LungLNP/RNA-1 did not further improve tumor control in this experiment, indicating that RNA-1 reached maximal therapeutic potential under the examined conditions (**Fig. 3b,h,i**). Across all groups, body weight and clinical tolerability indication were stable (**Supplementary Fig. 8**), and survival tracked with tumor growth trends. Mice treated with PBS, anti-PD-1, or LiverLNP/RNA-1 showed median survival times of approximately 15, 14 and 16 days after dosing, respectively, whereas treatment with LungLNP/RNA-1 prolonged survival to 25 days, with a comparable outcome (24 days) observed for the combination with anti-PD-1 (**Fig. 3c**). These results importantly show that the anticancer efficacy observed with LungLNP/RNA-1 correlates with the effect of this treatment on PD activity (increased levels of circulating interferons/cytokines) whereas treatment with LiverLNP/RNA-1 results in neither cancer efficacy nor PD effect.

**Fig. 3.**
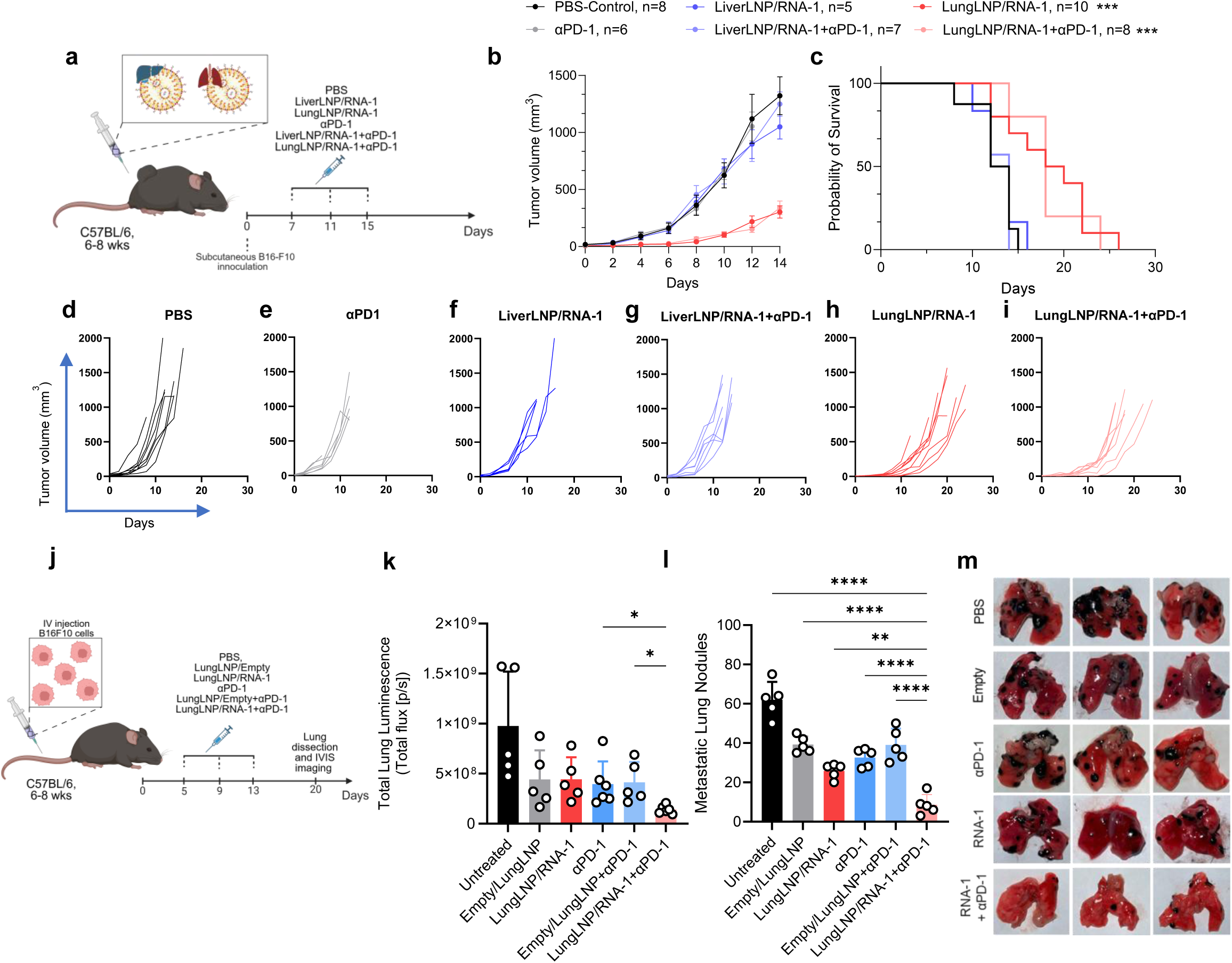
LungLNPs/RNA-1 suppresses tumor growth in subcutaneous and lung metastatic models of melanoma and demonstrates an additive response with immune checkpoint blockade in lung metastatic model. (a) Schematic of therapeutic regimen in the subcutaneous B16-F10 melanoma model. Mice were inoculated on day 0 and dosed intravenously with LungLNP/RNA-1 or LiverLNP/RNA-1 (2.2 mg/kg RNA-1) or control formulations, with or without intraperitoneal (i.p) aPD-1 antibody (100 μg), administered on days 7, 11, and 15 post tumor inoculation. (b) Average tumor growth curves in mice. (c) Kaplan–Meier survival curves for experimental cohorts. (d-i) Individual tumor growth curves for each treatment group. (j) Experimental outline of lung metastatic model followed by treatment with RNA-1 formulations and/or aPD-1. (k-m) Quantification of metastatic burden in lungs by total cell luminescence using IVIS imaging (k), nodule count (l), and representative lung images (m) demonstrating marked suppression of lung metastases in LungLNP/RNA-1 in combination with aPD-1. Data are represented as mean ± SEM from a representative experiment of two independent experiments with n = 5–10 (A–I) and mean ± SD from a representative experiment of two independent experiments, n = 5–6 (j–n) biologically independent samples. The data were analyzed by ordinary one-way ANOVA with Tukey’s multiple-comparisons test (b, c, l, m) and by Kruskal–Wallis test for pre-specified comparisons of RNA-1 vs combination and αPD-1 vs combination. Panels a and j were created with BioRender.com. * *P* ≤ 0.05, ** *P* ≤ 0.01, *** *P* ≤ 0.001, **** *P* ≤ 0.0001.

Realizing the potential role of the lungs in mediating RNA-1-driven immune activation, we also evaluated LungLNP/RNA-1 in a B16-F10 lung-metastasis model with or without anti-PD-1. C57BL/6 mice were inoculated intravenously with luciferase-expressing B16-F10 cells and treatment was initiated five days after injection. Lungs were collected on day 20 after inoculation, guided by the clinical status of control PBS-treated mice (**Fig. 3j**). Results based on tumor luminescence demonstrate that all treatments (anti-PD-1, LungLNP/RNA-1, and LungLNP/Empty with and without anti-PD-1) responded the same as control PBS treatment with significant tumor luminescent signal detected in the lungs of these animals (**Fig. 3k**). However, combination of LungLNP/RNA-1 with anti-PD1 reduced lung-associated tumor luminescence significantly. In addition, we observed an effect on metastatic nodule counts, with only the combination of LungLNP/RNA-1 and anti-PD-1 reducing the number of metastatic nodules significantly (**Fig. 3l**). Lung images supported these findings, showing few tumors in the combination-therapy group (**Fig. 3m**). Taken together, these findings clearly demonstrate an additive effect of innate immune agonism with immune checkpoint blockade in this aggressive and challenging to treat experimental lung metastasis model^64,65^ (**Fig. 3l,m**).

### RNA-1 activates innate and adaptive immunity

To characterize the immune phenotypes associated with LNP-enabled RNA-1 delivery, we profiled the myeloid and lymphoid lineages in B16-F10 mice with subcutaneous tumors after the administration of LiverLNP/RNA-1, LungLNP/RNA-1, or empty-LNP controls. In the spleen, RNA-1 in both Lung-and LiverLNPs increased the activation of dendritic cells (CD86), the activation and proliferation of CD8^+^ T cells, and the activation of CD4^+^ T cells and NK cells (**Fig. 4a(i); Supplementary Fig. 9a–d**), consistent with systemic immune engagement. LungLNP/RNA-1 selectively enhanced macrophage repolarization into a pro-inflammatory state (increased CD86/CD206 ratio) (**Fig. 4a(ii)**) and increased the activation of the cross-presenting dendritic cell population, including enhancement of CD86 expression in CD103⁺ DCs and CD8⁺ DCs, compared with LiverLNPs (**Fig. 4a(iii)–(iv)**).

**Fig. 4.**
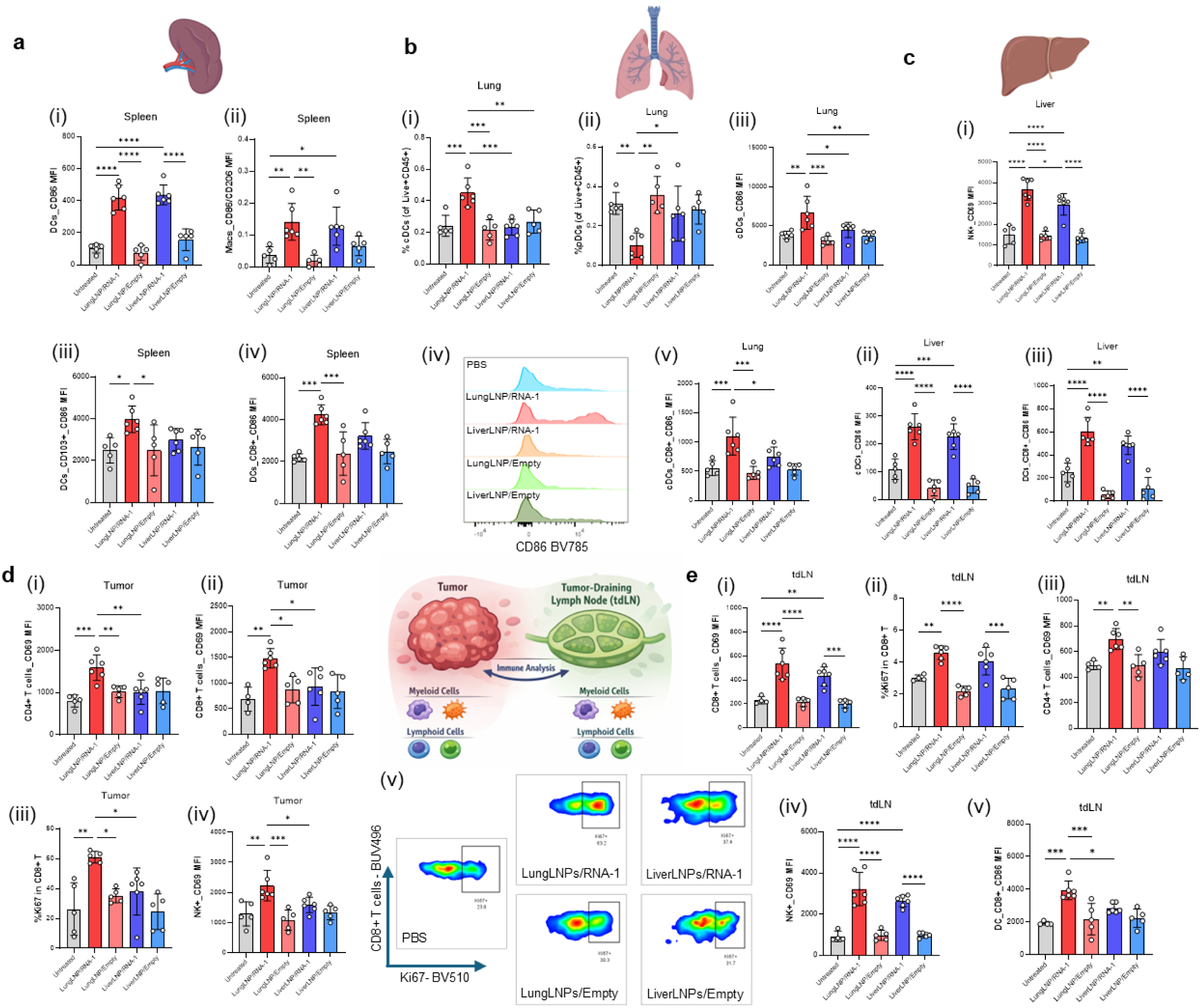
LNP organ BD licenses productive antitumour immunity of RNA-1 through differential early innate and adaptive immune activation. Immune phenotyping performed in B16-F10 subcutaneous tumor bearing mice and following systemic administration of RNA-1 using LungLNPs or LiverLNPs and corresponding vehicle controls and PBS. Immune populations and activation were analyzed using flow cytometry of the spleen, lung, liver, tumor microenvironment (TME) and tumor draining lymph node (tdLN), at two days post treatment. (a) Analysis of the spleen depicting quantification of (i) CD86 mean fluorescence intensity (MFI) in DCs (F4/80^-^/CD11c^+^/Ly6G^-^/CD45^+^), (ii) CD86/CD206 mean fluorescence intensity in macrophages (F4/80^+/^CD11b^+^/Ly6G^-^/CD45^+^), (iii) CD86 MFI in CD103^+^ DCs (CD103^+^/F4/80^-^/CD11c^+^/Ly6G^-^/CD45^+^), (iv) CD86 MFI in CD8^+^ DCs (CD8^+^/F4/80^-/^CD11c^+^/Ly6G^-^/CD45^+^). (b) Analysis of the lungs depicting quantification of (i) percent of DCs out of Live CD45^+^ cells, (ii) percent of pDCs out of Live CD45^+^ cells (CD11c^+^/CD24^-^/CD64^-^/SSC^hi^/CD11c^+^/CD11b^+^/Ly6G^-^/CD45^+^), (iii) CD86 MFI in DCs and (iv) representative histograms, (v) CD86 MFI in CD8^+^ DCs. (c) Analysis of the liver depicting quantification of (i) CD69 MFI in NK cells (NK1.1^+^/B220^-^/CD3^-^/CD11c^-^CD11b^-^/Ly6G^-^/CD45^+^), (ii) CD86 MFI in DCs (iii) CD86 MFI in CD8^+^ DCs. (d) Analysis of the TME depicting quantification of (i) CD69 MFI in CD4^+^ T cells (CD4^+^/NK1.1^-^/B220^-^/CD3^+^/CD11c^-^CD11b^-^/Ly6G^-^/CD45^+^), (ii) CD69 MFI in CD8^+^ T cells (CD8^+^/NK1.1^-^/B220^-^/CD3^+^/CD11c^-^CD11b^-^/Ly6G^-^/CD45^+^), (iii) precent of Ki67 positive CD8^+^ T cells. (iv) CD69 MFI in NK cells, (v) representative dot-plots showing the percent of Ki67^+^ CD8^+^ T cells. (e) Analysis of the tdLN depicting quantification of (i) CD69 MFI in CD8^+^ T cells, (ii) percent of Ki67^+^ CD8^+^ T cells, (iii) CD69 MFI in CD4^+^ T cells, (iv) CD69 MFI in NK cells, (v) CD86 MFI in CD8^+^ DCs. Data are represented as mean ± SD with n = 5-6 biologically independent samples. The data were analyzed by ordinary one-way ANOVA with Tukey’s multiple-comparisons test. * *P* ≤ 0.05, ** *P* ≤ 0.01, *** *P* ≤ 0.001, **** *P* ≤ 0.0001.

In the lung, LungLNP/RNA-1 increased the fraction of dendritic cells among lung CD45^+^ cells (**Fig. 4b(i)**), whereas in the liver, no such enrichment was evident under any condition (**Supplementary Fig. 10**). LungLNP/RNA-1 also reduced the percentage of plasmacytoid dendritic cells in the lung (**Fig. 4b(ii)**), consistent with RIG-I and TLR7 activation^66–68^. Moreover, delivery of RNA-1 using LungLNPs upregulated CD86 on lung dendritic cells (**Fig. 4b(iii)–(iv)**) and selectively increased CD86 expression on CD8⁺ DCs (**Fig. 4b(v)**). In the liver, induction of markers of NK cell activation (CD69) and DC activation (CD86), including on CD8^+^ DCs, were comparable between LungLNP/RNA-1 and LiverLNP/RNA-1, indicating that RNA-1 engages liver-resident antigen-presenting cells to similar extents with both formulations in the liver (**Fig. 4c(i)–(iii)**).

Within the tumor microenvironment, intratumoral dendritic cells and macrophages showed limited activation across all groups (**Supplementary Fig. 11a,b**). However, LungLNP/RNA-1 selectively induced the activation and proliferation of effector lymphocytes in tumors. RNA-1 delivered using LungLNPs also increased NK cell activation (CD69 expression) and proliferation (Ki67⁺ NK cells), and increased CD8^+^ T cell activation and proliferation (**Fig. 4d(i)–(v); Supplementary Fig. 11c**), along with CD4^+^ T cell activation (**Fig. 4d(iv)**). In the tdLNs, both formulations activated and induced proliferation of CD8^+^ T cells and activated NK cells (**Fig. 4e(i)–(iii)**), demonstrating that LungLNP/RNA-1 and LiverLNP/RNA-1 both trigger immune responses in tdLNs. However, the former more strongly increased activation markers on antigen-presenting cells, including CD86 expression on DCs (**Supplementary Fig. 12a**), CD8⁺ DCs (**Fig. 4e(iv)**), and macrophages (**Supplementary Fig. 12b**), as well as selectively activating CD4^+^ T cells.

Together, these findings demonstrate that RNA-1 drives lymphoid and myeloid immune activation, with lung-specific delivery enabled by LungLNPs shaping the magnitude and quality of the response as well as the cell types engaged. LungLNP delivery amplifies cross-presenting dendritic cell activation and effector lymphocyte proliferation, highlighting how organ-selective LNP distribution dictates downstream immune responses.

### LungLNP/RNA-1 suppress human tumor growth in a human Lung Cancer Chip

To assess the translational potential of RNA-1-mediated innate immune activation in a human-relevant lung tumor microenvironment, we modified a previously described human Lung Cancer Chip microfluidic culture model that recapitulates key structural and functional features of the human alveolar niche and organ-specific growth patterns observed in human patients^69–71^. Lung Cancer Chips containing GFP-labeled A549 lung cancer cells grown within a healthy human alveolar epithelium interfaced with a pulmonary microvascular endothelium within a two-channel Organ Chip device (**Fig. 5a**) were treated with either LungLNPs encapsulating RNA-1 at 100 nM or 200 nM, or a matched empty-LNP control (200 nM), following two doses of the formulation being perfused through the vascular channel (to mimic intravenous administration) every 4 days as in the mouse studies (**Fig. 5b**). Fluorescence microscope imaging of A549 GFP fluorescence intensity over time revealed a pronounced, dose-dependent suppression of human lung cancer cell growth in LungLNP//RNA-1-treated chips compared with untreated chips and chips treated with empty LNPs (**Fig. 5c**). Tumor growth inhibition became evident following the second dosing cycle and was most pronounced at the 200 nM dose. Moreover, representative fluorescence imaging on day 11 revealed a substantial reduction in A549 cell density and spatial coverage in the 200 nM LungLNP/RNA-1 treated group in comparison to the untreated control group, which exhibited dense tumor growth filling the surface of the entire epithelial channel (**Fig. 5d**). These findings demonstrate that RNA-1 can inhibit the growth of human tumors in a physiologically relevant organotypic microenvironment, as well as inhibit mouse tumors.

**Fig. 5.**
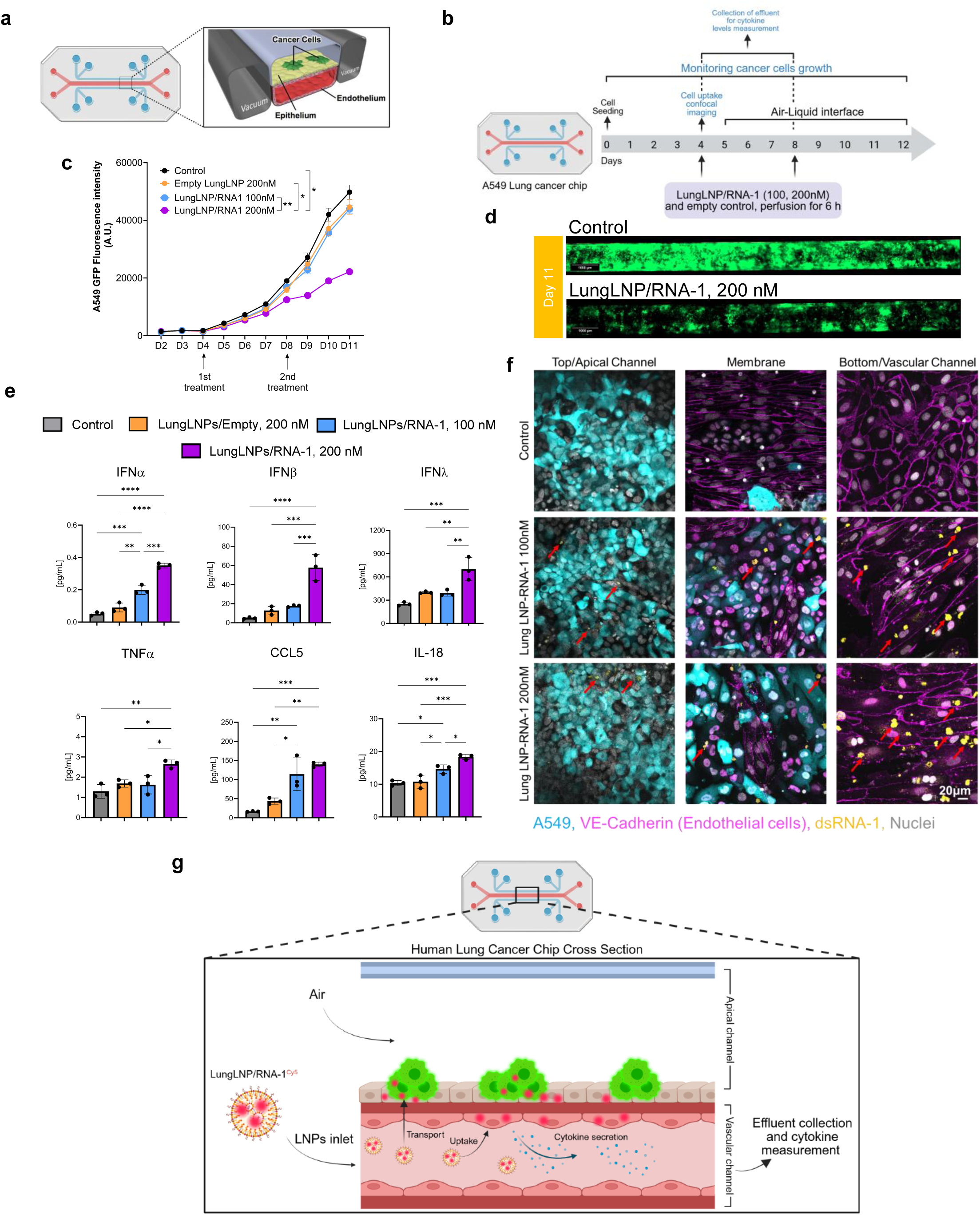
RNA-1 suppresses tumor growth and interacts with the endothelium in a human Lung Cancer Chip model. (a) Schematic illustration depicting a cross-section of the human lung cancer chip model, which recapitulates key physiological and pathophysiological features of human lung cancer. The microfluidic chip top channel containing human lung epithelial cells and human A549 adenocarcinoma alveolar basal epithelial cells stably expressing GFP, bottom channel containing human lung microvascular endothelial cells cultured on all four walls of the lower channel. (b) Treatment regimen for the human lung cancer-chip using LungLNPs/RNA-1 (100 and 200 nM), and empty LungLNP control (LungLNPs/Empty, 200 nM) and untreated chips. The first treatment was administered 4 days post-seeding, followed by establishment of the air–liquid interface on the same day. A second dose was administered on day 8. LNPs were delivered by vascular perfusion for 6 h per treatment. (c) A549 tumor growth curves during the treatment regimen, quantified by longitudinal GFP fluorescence imaging and measurement of fluorescence intensity. Data were analyzed using a two-way mixed effects model with time and treatment as fixed effects, followed by Tukey’s multiple-comparisons test. (d) Representative fluorescence images showing A549 tumor cells (green) on day 11 (scale bar = 1000 µm). (e) Quantification of cytokines and chemokines measured 2 h following the second dose. Data were analyzed by one way ANOVA with Tukey’s multiple comparisons test. (f) LNP uptake in the lung cancer-chip following perfusion of fluorescently labeled LungLNPs/RNA-1^Cy5^ (yellow) at 100 and 200 nM. Endothelial cells were stained for VE-cadherin (purple), A549 tumor cells expressing GFP are shown in blue, and nuclei are shown in white. Chips were imaged 4 days post-treatment using confocal microscopy (scale bar = 20 µm). (g) Schematics depicting the mechanistic insight into RIG I-mediated lung cancer immunotherapy in human lung cancer chip demonstrating internalization into endothelial cells and RIG I activation and secretion of cytokines. Uptake into epithelial cells via direct exposure or via transport through gaps in the endothelial barrier. Panels a, b and d were created with BioRender.com. * *P* ≤ 0.05, ** *P* ≤ 0.01, *** *P* ≤ 0.001, **** *P* ≤ 0.0001.

To profile cytokine responses within the tissues in the chip, we collected vascular effluent and measured cytokines and chemokines 2 h after the first (**Supplementary Fig. 13**) and second dosing (**Fig. 6e**). LungLNP/RNA-1 increased type I and III IFNs, inflammatory cytokines, and chemokines in a dose-dependent manner, with the strongest responses at 200 nM, which correlated with optimal tumor growth inhibition (**Fig. 5c**). IFN *α* increased after both the first and second doses (5.2-and 8-fold, respectively), whereas IFN *β* increased strongly after the first dose (26.7-fold) and remained elevated after the second dose (11.7-fold). LungLNP/RNA-1 also increased IFN *λ*, CCL5, IL-18, TNF *α*, and IL-6, particularly after the second exposure (**Fig. 6e**).

Finally, to assess LNP transport and cellular uptake in the chip, we perfused fluorescently labeled LungLNP/RNA-1 through the vascular channel at 100 nM and 200 nM. Confocal imaging detected Cy5 signal in multiple compartments, with prominent uptake in VE-cadherin^+^ endothelial cells lining the vascular channel as well as detectable Cy5 signal in both the alveolar epithelial cells and A549 tumor cells (**Fig. 6f**). Uptake was most prominent in endothelial cells, which is consistent with the preferential uptake into endothelial cells that we observed in the mouse BD data.

## Discussion

In this work, we have shown that changes in the tissue and cellular exposure to an immunostimulatory duplex RNA, imparted by use of different delivery vehicles, can gate the in vivo PD and antitumor activity of its downstream immunomodulatory responses, even when in vitro potency is similar across delivery formulations. Specifically, we used two LNP formulations with comparable physical properties and equivalent in vitro potency, differing in lung and liver tropism to deliver RNA-1. We observed a marked discrepancy in systemic cytokine PD, with the LungLNP formulation producing robust circulating IFNs and inflammatory cytokines, whereas the levels elicited by the liver-biased formulation remained near those of the empty LNP controls. We found a similar pattern when using the lung-tropic polymeric carrier JetPEI, which supports pulmonary tropism as a major contributor to augmented systemic cytokine responses. Interestingly, this reliance on pulmonary delivery differs from other RIG-I agonists such as stem-loop RNA molecules^12,72^ or MK-4621^73,74^ that reached clinical trials, or TLR7 agonists^75,76^.

Beyond shifting organ distribution, the LungLNP formulation also enabled the distribution to and activation of specific pulmonary cell populations. Indeed, LungLNPs induced more pronounced DC activation in the lungs compared to LiverLNPs with a notable reduction in pDCs, possibly due to a negative feedback mechanism in response to prolific IFN-I production. Across BD, tissue cytokine measurements and immune phenotyping, our data are consistent with pulmonary endothelial, epithelial, and immune cell populations acting as a specialized sensing network for non-self RNA that amplify the systemic cytokine response and enable robust anticancer efficacy^77–81^. These observations are consistent with recent evidence that the lung deploys a spatially tiered threat-sensing mechanism, in which in relationship to external viral exposure, deeper stromal compartments are more poised than the outer epithelia to initiate innate immune responses^24^. Viewed in this context while administering RNA-1 systemically via LNPs, lung-tropic delivery may enhance RNA-1 activity not only by changing organ exposure, but also by increasing access to pulmonary cell populations with higher intrinsic capacity such as endothelial cells to sense and respond to immunostimulatory RNA.

More specifically, the differences in cytokine PD across LNP formulations correlated with differences in efficacy, in particular, enhanced antitumor activity with the LungLNP formulation in the subcutaneous melanoma model, and additivity with anti-PD-1 in the melanoma lung metastasis model. These correlations are consistent with the magnitude and timing of systemic cytokine secretion and exposure influencing downstream immune phenotypes and therapeutic outcomes^82^. Accordingly, we found that only treatment with LungLNPs activated lymphocyte populations and NK cells in the tumor, likely due to systemic cytokine production but potentially contributed to by endothelial cell-immune cell crosstalk as we found LungLNPs in tumor-associated endothelial cells to a greater extent than LiverLNP treated mice.

Notably, both LungLNP/RNA-1 and LiverLNP/RNA-1 formulations showed evidence of engaging local innate pathways outside the lung. Liver and spleen homogenates showed similar IFNα levels between formulations early after dosing, and both formulations induced comparable immune activation phenotypes in the spleen and liver. However, only LungLNP/RNA-1 exhibited additional activity in the lungs that were correlated with therapeutic efficacy, even when tumors were grown subcutaneously. Together, these results highlight that shifting delivery of immunostimulatory RNA-1 to the lungs is crucial for translating innate immune activation into effective antitumor responses even at distant sites.

The murine knockout data further revealed which innate pathways contribute to the measured plasma cytokine phenotype. The response was strongly reduced in TLR7-deficient mice and also in mice lacking RIG-I although to a lesser degree, which is consistent with combined endosomal and cytosolic sensing. This contrasts with exclusive RIG-I-dependent IRF3 activation demonstrated in cultured epithelial cells treated with RNA-1 with an in vitro transfection agent^25^, which underscores that receptor usage can vary with tissue context and/or with different delivery formulations. The potential clinical relevance of our findings is further reinforced by results from the human Lung Cancer Chip study, which demonstrated robust IFN and pro-inflammatory cytokine secretion in response to administration of LungLNP/RNA-1. Importantly, RNA-1 delivered with LungLNP induced tumor suppression, and its preferential uptake into endothelial cells underscores their potential contribution to tumor control.

Several considerations should be considered when interpreting the delivery-PD relationships identified in this study. BD and cellular uptake were assessed using fluorescently labeled RNA, which reports localization but does not directly quantify the exposure to intact RNA-1; complementary analytical assays measuring dsRNA-1 levels will be necessary further strengthen exposure-response analyses in the future. In addition, while we characterized the organs and cell populations exposed to RNA-1, we did not resolve the cellular sources responsible for initiating the early cytokine response. More broadly, the immune outcomes observed here may be specific to the combination of RNA-1 and LungLNP delivery, and testing additional innate immune agonists delivered to different organs may reveal distinct cytokine responses depending on the tissue and cellular target of delivery. In the human Lung Cancer Chip model, the LungLNP/RNA-1 increased cytokine secretion and reduced the growth of A549 cells; however, because the chip lacks immune cells, these experiments demonstrate tissue-intrinsic responses that appear to significantly impact tumor growth locally, but they do not model immune-mediated tumor control. Finally, although lung-tropic LNP formulations are currently being evaluated in several clinical trials, none have yet received clinical approval, highlighting that further understanding of the impact of modifying LNP chemistry is still important for effective clinical translation of this organ-specific approach. Collectively, our findings demonstrate that organ and cellular context must be treated as fundamental design constraints in the development of next-generation delivery vehicles for immunomodulatory RNA therapeutics.

## Acknowledgements

This research was, in part, funded by the Advanced Research Projects Agency for Health (ARPA-H). The views and conclusions contained in this document are those of the authors and should not be interpreted as representing the official policies, either expressed or implied, of the United States Government.

We thank Wontaek Chung for his assistance and support with in vivo mouse experiments at the Koch Institute for Integrative Cancer Research at the Massachusetts Institute of Technology (MIT); the Wyss Institute veterinary team for their support and assistance with in vivo experiments; and Eric Zigon from the Wyss institute’s flow cytometry core for consultation on flow cytometry panel design and support in experiments. Biorender was used to create schematics.

## Contributions

E. A-L., A.M.C., K.E.C., D.E.I. and N.A. conceptualized the study and wrote or edited the original drafts of the manuscript. N.A.I., W.S., Z.P., N.R., J.C.O.E., N.N.C.H., S.G.B., E. A-L. and A.M.C. performed *in vitro* studies or *ex vivo* processing. J.J., C.B., J.F.K.S. and Y.M. performed human chip studies. A.G., M.S.S., E. A-L. and A.M.C. performed *in vivo* studies. A.E. and W.M.S. provided materials and computational analysis. G.G., K.E.C., W.M.S., D.E.I. and N.A. supervised the study and were responsible for funding acquisition.

## Methods

***RNA-1 structure*** RNA-1 was custom synthesized by IDT with the following sequence, sense: 5’-CUGAUGACACUGGCUAGUUCACCTT-3’, antisense:

GGGACUACUGUGACCGAUCAAGUGGAA. Sense and antisense RNA strands were annealed in nuclease-free Duplex Buffer (Integrated DNA Technologies, 11-05-01-03) using a T100 thermal cycler (Bio-Rad) with the following program: 90 °C for 1 min, 75 °C for 1 min, followed by slow cooling at 1 °C per 1.5 min.

***Fabrication of LNP formulations*** LNP formulations were prepared by pipette mixing of an organic lipid phase with an aqueous RNA phase at a 1:3 (v/v) organic:aqueous ratio. The organic phase consisted of a lipid mixture dissolved in ethanol, while the aqueous phase contained RNA-1 diluted in 10 mM sodium acetate buffer (pH 5.2; Sigma, 567422). The lipid mixture included the ionizable lipid SM-102 (BroadPharm, BP-25499), helper lipid 1,2-dioleoyl-sn-glycero-3-phosphoethanolamine (DOPE; Avanti Polar Lipids, 850725P), cholesterol (Sigma, C8667), and 1,2-dimyristoyl-rac-glycero-3-methoxypolyethylene glycol-2000 (DMG-PEG2000; Avanti Polar Lipids, 880151P), at a total lipid to RNA mass ratio of 20:1. LiverLNPs/RNA-1 were formulated at a molar ratio of 50:10.5:38:1.5 (SM-102/DOPE/cholesterol/DMG-PEG2000). LungLNPs/RNA-1 (LungLNPs) included an additional permanently charged lipid, 1,2-dioleoyl-3-trimethylammonium-propane (DOTAP; Avanti Polar Lipids, 890890P), at a molar ratio of 25:5.3:19:0.8:50 (SM-102/DOPE/cholesterol/DMG-PEG2000/DOTAP). Following assembly, LNPs were dialyzed against phosphate buffered saline (PBS; pH 7.4) using 3.5 kDa molecular weight cut-off dialysis tubes (Sigma, PURD35050) to remove residual ethanol prior to downstream characterization and use

***Characterization of LNPs formulations*** Particle size and polydispersity indices of LNP formulations were measured by dynamic light scattering (DLS) using a Zetasizer Nano ZS instrument (Malvern Instruments, 633 nm laser) at a backscattering angle of 173°. Surface charge measurements were obtained by phase analysis light scattering. Measurements were performed on freshly prepared samples, diluted in PBS, at room temperature. Encapsulation efficiency of RNA-1 within LNPs was quantified using the modified Quant-iT RiboGreen RNA assay (Invitrogen). LNP samples were diluted in either TE buffer (to measure free, unencapsulated RNA) or TE containing 2% Triton X-100 (to fully disrupt particles and release encapsulated RNA). Duplicates of LNPs and standard curve samples (100 μL) were dispensed into a black 96-well microplate, followed by an addition of 100 μL of RiboGreen reagent (1:200 dilution in TE). Fluorescence was recorded using a TECAN Infinite M Plex plate reader (λex=485 nm, λem=528 nm). The fluorescence intensity from intact LNPs (I_intact) corresponded to free RNA, while the intensity after Triton disruption (I_disrupted) represented total RNA content. Encapsulation efficiency (%EE) was calculated as: %EE = 100 × (1-(Iintact/Idisrupted))^1^.

***Cell culture and isolation of bone marrow derived dendritic cells (BMDCs)*** *Mus musculus* B16-F10 melanoma and B16-F10-Luc (ATCC) were used to inoculate mice *in vivo*. Human NF-κB-SEAP & IRF-Luc Reporter lung carcinoma (A549) cells (A549 RIG I) and human RIG-I-KO Dual Reporter A549 cells (A549 RIG I KO) (InvivoGen) were used to study the in vitro innate immune activity of RNA-1. All cell lines were maintained in Dulbecco’s Modified Essential Medium (DMEM, Gibco, no.11995-065) with D-glucose and L-glutamine supplemented with 10% (v/v) heat inactivated fetal bovine serum (HI-FBS, Gibco, no.10500-064), and 1% (v/v) penicillin-streptomycin (Pen Strep, Gibco, no. 15140-122). Selection antibiotics were applied as follows: A549 cultures were maintained with the addition of Zeocin™ (InvivoGen, ant-zn-05; 100 µg/mL) and Blasticidin™ (InvivoGen, ant-bl-1; 10 µg/mL) at every other passage, and B16-F10-Luc cultures were maintained with blasticidin (10 µg/mL). All cells were cultured at 37 °C in a humidified incubator with 5% CO₂.

Bone marrow derived dendritic cells (BMDCs) were isolated as described before^2^. Briefly, bone marrow was flushed from the femurs and tibias of 6-8-week old female wild-type C57BL/6 mice using ice cold HBSS (Gibco, no. 14025092) and passed through a 40 μm cell strainer. Bone marrow stem cells were pelleted and resuspended in ACK lysis buffer (Gibco, no. A1049201) for 1 min and washed with HBSS. Afterwards, cells were seeded using complete medium (RPMI-1640, 10% (v/v) HI-FBS, 1% (v/v) penicillin/streptomycin) in petri dishes supplemented with recombinant murine granulocyte-macrophage colony-stimulating factor (GM–CSF, BioLegend, 20 ng/mL). The culture medium containing GM-CSF was replaced on days 3, 6, and 8.

### In vitro assays

***Assessment of RIG-I activity via IRF3 reporter cells*** Cells were seeded at 1.5 x 10^4^ cell per well in a 96-well plate 24 h prior to treatment. Subsequently, A549 RIG I and A549 RIGI KO cells were treated with Liver or LungLNPs/RNA-1 or their empty counterparts in 120 μL and 48 h post-treatment, IRF3 reporter activity was assessed using the Quanti-Luc (Invivogen) assay per the manufacturer’s instructions. Briefly, 50 μL reagent into a white plate preloaded with 20 μL supernatant and luminescence was immediately read on a Synergy H1 multimodal microplate reader. Subsequently, NFκb activity was measured using Quanti-Blue (InvivoGen) by adding 180 μL reagent to 20 μL supernatant in a clear plate, incubating for at least 30 min at 37°C, and absorbance was read on the Synergy H1 microplate reader. Cell viability was assessed with CellTiter-Glo® (Promega) by adding reagent 1:1 (v/v) to the residual medium in each well, incubating 10 min to stabilize signal, transferring to a white plate, and recording luminescence.

***BMDC activation*** Cells were seeded at 1 x 10^5^ cells per well in a 24-well plate 24 h prior to treatment. Subsequently, BMDCs were treated with Liver or Lung LNPs/RNA-1 or their empty LNPs control in 450 μL at a final concentration of 10 μM. At 24 h post-treatment the cells were detached, washed with cell staining buffer (CSB, BioLegend) and stained using fluorescently labeled anti-mouse antibodies against CD11c (BV421, N418 clone), MHC-II (BV605, M5/114.15.2 clone), CD80 (FITC, 16-10A1 clone), CD86 (PE, PO3 clone), and CD40 (APC, 3/23 clone). To exclude dead cells, LIVE/DEAD NIR (Thermo Fisher) was used. Following staining, cells were fixed using fixation buffer (BioLegend) and washed before loading into a 96-well plate U-bottom (Eppendorf). Stained cells were analyzed using flow cytometry with a Sony iD7000 spectral flow cytometer and the data were analyzed using FlowJo software.

***Animal care and experimentation*** Wild-type (WT) female C57BL/6 mice (6-10 weeks old) were obtained from Charles River Laboratories. Female mice (6-10 weeks old) lacking key RNA sensing mechanisms, RIG-I (C57BL/6NJ-*Rigi^em1(IMPC)J^*/Mmjax, #046070, RIG-I KO), TLR3 (B6;129S1-*Tlr3^tm1Flv^*/J, #005217, TLR3 KO) and TLR7 (B6.129S1-*Tlr7^tm1Flv^*/J, #008380, TLR7 KO) were obtained from Jackson Labs. All animal research and veterinary care were conducted in compliance with protocols approved by the Institutional Animal Care and Use Committee (IACUC) at the Koch Institute for Integrative Cancer Research at the Massachusetts Institute of Technology (MIT) and the Harvard Medical Area Office of the IACUC (OOTI). Mice were housed under a 12-hour light/dark cycle and had access to water and standard rodent diet (LabDiet 5053, irradiated) ad libitum.

***Mouse pharmacodynamic (PD) model experiments*** WT female C57BL/6 mice (6–8 weeks old) or RIG-I/TLR3/TLR7 KO mice (6-8 weeks) were administered intravenously (i.v.) with dsRNA-1 encapsulated in either LungLNP or LiverLNP formulations (2.2 mg/kg in 100 µL). Control mice were treated with PBS or empty LNPs corresponding to each organ-tropic formulation. JetPEI was included as a positive control for the LungLNPs in which nanoparticles were administered either i.v. or intraperitoneally (i.p.). Blood samples were collected via the submandibular route at 2 and 6 h, and post-mortem cardiocentesis at 24 h post-administration and immediately placed on ice. The samples were centrifuged at 2500 g for 15 minutes at 4 °C to separate the plasma, which was subsequently analyzed for IFNα, IFNβ, IFNγ and TNFα (MSD assay, K15320K-2) and IFNλ (IL-28A, Abcam, ab208989).

***AlphaFold3 modeling*** AlphaFold3 was used to model how RNA-1 could engage mouse and human RNA-sensing receptors. Protein sequences for mouse and human RIG-I and TLR7 (UniProt entries O95786, Q6Q899, Q9NYK1, P58681) along with RNA-1 strand sequences were submitted to the AlphaFold Server using default settings. To explore potential multivalent assemblies, complex predictions were run for (i) four copies of RIG-I with two copies of each RNA-1 strand and (ii) two copies of TLR7 with single strands of RNA-1. The highest-ranked predictions were inspected to assess plausible receptor–RNA binding modes and rendered using UCSF ChimeraX (v1.7).

***Efficacy study in B16-F10 subcutaneous tumor model*** B16-F10 subcutaneous tumors were established by injection of 0.5×10^6^ cells (suspended in 50 µL of cold sterile HBSS) subcutaneously into the lower right flank of female C57BL/6 (6-8 weeks old) as described before^3^. Treatment regimen included i.v. administration of three doses of PBS (100 µL), dsRNA-1 (2.2 mg/kg in 100 µL) encapsulated in either Lung or Liver-tropic LNPs or corresponding empty LNPs (2.2 mg/kg in 100 µL). Doses were administered on day 7, 11 and 15 post tumor inoculation. Mice that received dual treatment of both RNA-1 and anti-PD-1 (clone RMP1-14, Bio X Cell) dosed intraperitoneally (100 µg) on days 8, 12 and 16 post tumor inoculation. Tumor dimensions were measured and tumor volume was calculated using the following equation: V = L x W x H x π/6 and was measured along with body weight.

***Efficacy study in B16-F10 lung metastatic tumor model*** B16-F10 lung metastases were established by i.v. tail vein injection of 0.2×10^6^ luciferase expressing B16-F10 cells (B16-F10-Luc, suspended in 100 µL of cold sterile PBS) into female C57BL/6 mice (6-8 weeks old). Treatment regimen included i.v. administration of three doses of PBS (100 µL), RNA-1 (2.2 mg/kg in 100 µL) encapsulated in either Liver or LungLNPs or corresponding empty LNPs (2.2 mg/kg in 100 µL). Doses were administered on day 5, 9 and 13 post tumor inoculation. Mice that received dual treatment of both dsRNA-1 and immune checkpoint inhibitor therapy, were dosed intraperitoneally with 100 µg of aPD-1 (clone RMP1-14, Bio X Cell) on days 6, 10 and 14 post tumor injections. On day 22, mice were euthanized, and the lungs were carefully dissected, weighed, and placed in black 12-well plates (Cellvis)^4^. The tissues were then incubated in 1 mg/mL D-luciferin (IVISbrite D-Luciferin Potassium Salt, XenoLight) diluted in PBS. Ten minutes post incubation, bioluminescent images were acquired using IVIS imaging. Luminescence was quantified as total radiant flux (photons/second) for each lung. The lungs were also photographed, weighed, and examined to count the number of visible metastatic tumor nodules in each lobe.

***Organ-tropic LNPs BD*** Female C57BL/6 tumor bearing mice (6–8 weeks old) were administered i.v. with dsRNA-1 encapsulated in Lung-or Liver-tropic LNPs (2.2 mg/kg in 100 µL) in which 10% (w/w) of the dsRNA was labeled with Cy5. RNA-1 was labeled with Cy5 using a nucleic acid labeling kit (Mirus Bio™ Label IT™ Nucleic Acid Labeling Kit, MIR3725). Four h post dosing mice were perfused with PBS and euthanized. Organs (liver, lung, spleen, heart, kidneys, tumor and tdLN) were collected, immersed and washed with ice cold PBS and imaged using an IVIS Spectrum (Perkin Elmer) for the detection of dsRNA-1^Cy^^5^ fluorescence (λex= 640 nm, λem= 680 nm). Fluorescence intensities were calculated using Living Image software (Perkin Elmer). Following imaging, organs were weighed and frozen at-80 °C until further analysis.

***Organ cytokine levels*** Female C57BL/6 mice (6-8 weeks) bearing B16-F10 tumors were injected with one i.v. dose of PBS and RNA-1 encapsulated in LungLNP or LiverLNP at 2.2 mg/kg. After 2 h, mice were perfused with PBS and euthanized. Liver, lungs, spleen, tumor and tdLNs were dissected and blood was collected to obtain plasma. Tissues were homogenized (Precellys® CK14 Lysing Kit) and lysed using T-PER mild lysis buffer (78510, Thermo) containing 1:100 protease inhibitor cocktail (ab271306, abcam) and tissue homogenate were separated from debris. Levels of proinflammatory cytokines and chemokines IFNα, IFNβ, IFNγ, CXCL-1, CCL-2 (MCP-1), CCL5, CXCL-10, IL-1β, IL-6, TNFa IL-12, GM-CSF, IL-10 were measured using manufacturer’s instructions (Legendplex 740622, Biolegend).

***Immunophenotyping studies*** Tumor bearing female C57BL6/J mice (6–8 weeks old) were treated with PBS (100 µL), dsRNA-1 (2.2 mg/kg in 100 µL) encapsulated in either Lung or Liver-tropic LNPs or corresponding empty LNPs (2.2 mg/kg in 100 µL), all intravenously administered. Immune activation markers and immune lineage organ composition were analyzed 48 h post treatment. Lung, liver, spleen, tumor and tumor draining lymph node (tdLN) were collected from all mice. The liver, lungs and tumors were cut into small pieces and digested into single cells in 2 mL of HBSS solution (depleted of Ca2+ and Mg2+) containing type I collagenase (1 mg/mL, LS004214, Worthington Biochem) and DNase I (0.05 mg/ml, 10104159001, Sigma) in a 6 well plate that was then incubated at 37°C on an orbital shaker for 25 min. Tissues were extruded through an 18 G needle and returned to the shaker for additional 30 min. Following digestion, 3 mL of HBSS was added to stop the enzymatic activity. The digested tissues, spleen and tdLN were mashed and transferred through a 40 um cell strainer and washed. Pelleted cells from all tissues, except the tdLN, were treated with ACK Lysing Buffer (Gibco) to remove erythrocytes. The cells were washed with HBSS, passed through a 40 µm filter and counted. For staining, 1 × 10^6^ cells per sample were seeded and first labeled for surface markers, then fixed and permeabilized (562574, BD Biosciences), followed by permeabilization and intracellular staining. The list of anti-mouse antibodies, clones and dilutions that were used for flow cytometry are listed in Table S.1. Stained cells were analyzed by flow cytometry using a Sony iD7000 flow cytometer and all the data were analyzed using FlowJo software. Supplementary Fig S.14-16 show the gating strategy, adapted from Baumann *et al.*^5^.

***Human Lung Chip culture*** Human Lung Chips were cultured as previously described with minor modifications^6^. Briefly, two-channel Organ Chip (Chip S1, Emulate, Inc.) were activated and coated with collagen IV (200 μg/mL) and laminin (15 μg/mL). Human primary lung microvascular endothelial cells (Lonza, CC-2527; passage 6) were seeded into the vascular channel at a density of 8 × 10⁶ cells/mL. GFP-labeled A549 lung adenocarcinoma cells were seeded at a density of 1 × 10⁵ cells/mL and co-seeded with primary human alveolar epithelial cells (Cell Biologics, H-6621; passage 3) at a density of 5 × 10⁵ cells/mL into the apical channel. Chips were maintained under continuous perfusion (30 μL/h) with automatic fluid handling ZOE systems (Emulate, Inc.), using epithelial growth medium (Cell Biologics, H-6621) in the apical channel and EGM®-2MV microvascular endothelial growth medium (Lonza, CC-2527) in the vascular channel. Air-liquid interface (ALI) was established on day 4 post seeding, after which chips were perfused exclusively through the vascular channel using EGM®-2MV but with reduced fetal bovine serum (FBS) to 0.5% v/v, termed as modified EGM-2MV. Chips were cultured for an additional 3 days following ALI induction, after which the basal medium was switched to a modified EGM®-2MV formulation containing reduced hydrocortisone (50 nM instead of 550 nM in the standard formulation). dsRNA formulated in LungLNPs was administered by perfusion at a flow rate of 30 μL/min for 6 h on day 4 (both apical and vascular channels) and day 8 (vascular channel only) post seeding, after which the medium was replaced with fresh culture medium. Cancer cell growth was monitored by quantifying GFP fluorescence intensity from top-view images acquired every other day using an Axio fluorescence microscope (Zeiss). Image analysis was performed using ImageJ software. Regions of interest (ROIs) were manually selected to include the cell-populated co-culture area. The same ROI size and location were applied to all chips to ensure consistency. Fluorescence intensity was quantified as mean fluorescence intensity after background subtraction, using identical imaging settings for all samples.

***Cytokine analysis (Lung Chip)*** Effluents were collected from the basal or vascular channel of chips at 2 and 24h post treatment. Cytokine levels were quantified using a multiplex bead-based Luminex assay (custom ProcartaPlex panels, Thermo Fisher Scientific) according to the manufacturer’s instructions. Data was acquired on a Bio-Plex 3D system and analyzed using Bio-Plex Manager software (Bio-Rad).

***Confocal microscopy and image analysis (Lung Chip)*** On day 12 (experimental endpoint), chips were washed with DPBS (−/−), fixed with 4% paraformaldehyde for 10 min at room temperature, permeabilized with 0.2% Triton X-100 for 15 min, and blocked with 10% goat serum at 4 °C overnight. Chips were then incubated with a rabbit anti-VE-cadherin primary antibody (1:100; PA5-19612, Thermo Fisher Scientific) overnight at 4 °C, followed by incubation with an Alexa Fluor 555 conjugated goat anti-rabbit secondary antibody (1:1000; A-21428, Thermo Fisher Scientific) for 1 h at room temperature. Chips were then washed and mounted with 4′,6-diamidino-2-phenylindole (DAPI, D9542, Sigma Aldrich) before confocal microscopic imaging. Confocal imaging was performed using ZEISS LSM 980 microscopes.

***Statistical analysis*** Statistical analyses were performed using GraphPad Prism [11.0.0 (84)]. Data are presented as [mean ± s.d./mean ± s.e.m.], as indicated in the figure legends, which also define n and the number of independent experiments. In vitro experiments have a minimum of n=3 biological replicates. Pair-wise comparisons were done using two-tailed Welch’s t-test. Comparisons among multiple groups were performed using one-way or two-way ANOVA followed by Tukey’s multiple-comparisons test unless specifically specified in the legend. In vivo experiments were done using a minimum of n = 5 biological replicates per condition in each experiment.

## Supplementary Figures

**Supplementary Fig. 1.**
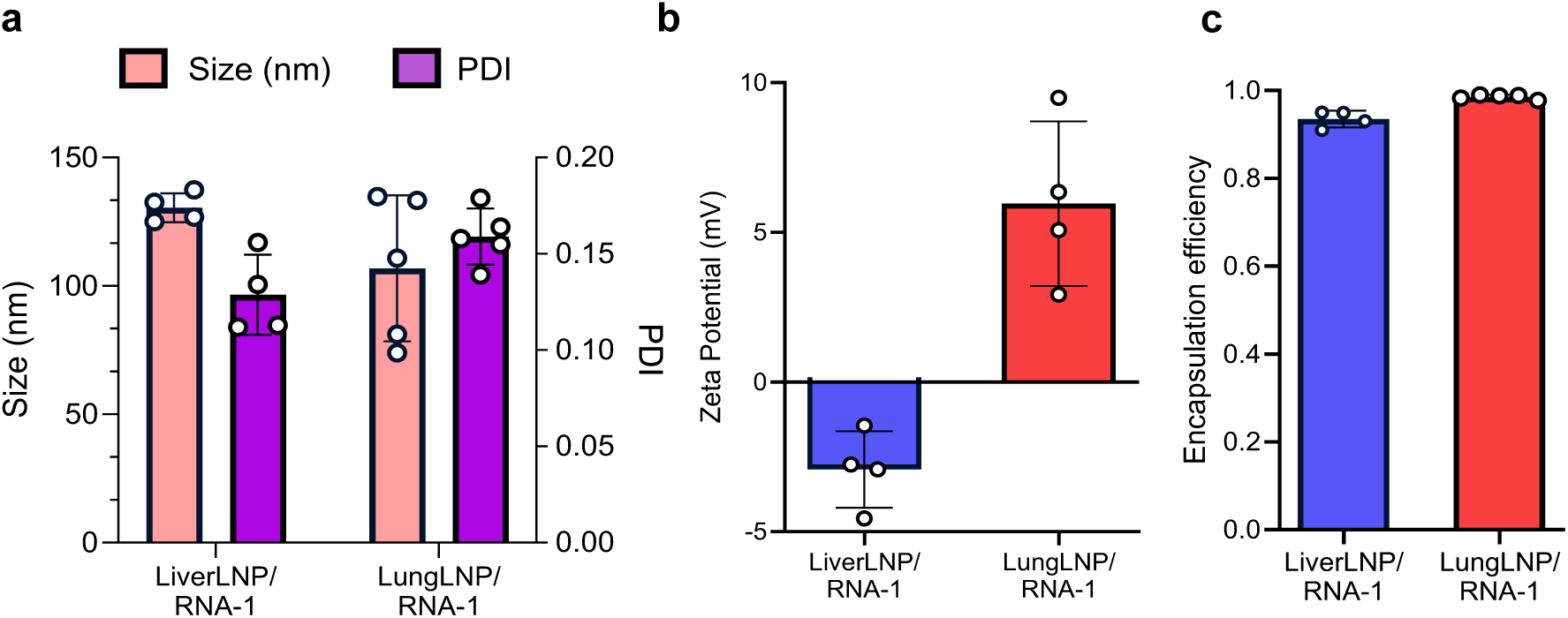
LiverLNP/RNA-1 and LungLNP/RNA-1 share comparable physical properties except for their surface charge. Physicochemical characterization of LiverLNP/RNA-1 and LungLNP/RNA-1 by (a) nanoparticle size and polydispersity by dynamic light scattering, (b) ζ-potential, and (c) encapsulation efficiency. Data presented as average ± SD, n = 4-5.

**Supplementary Fig. 2.**
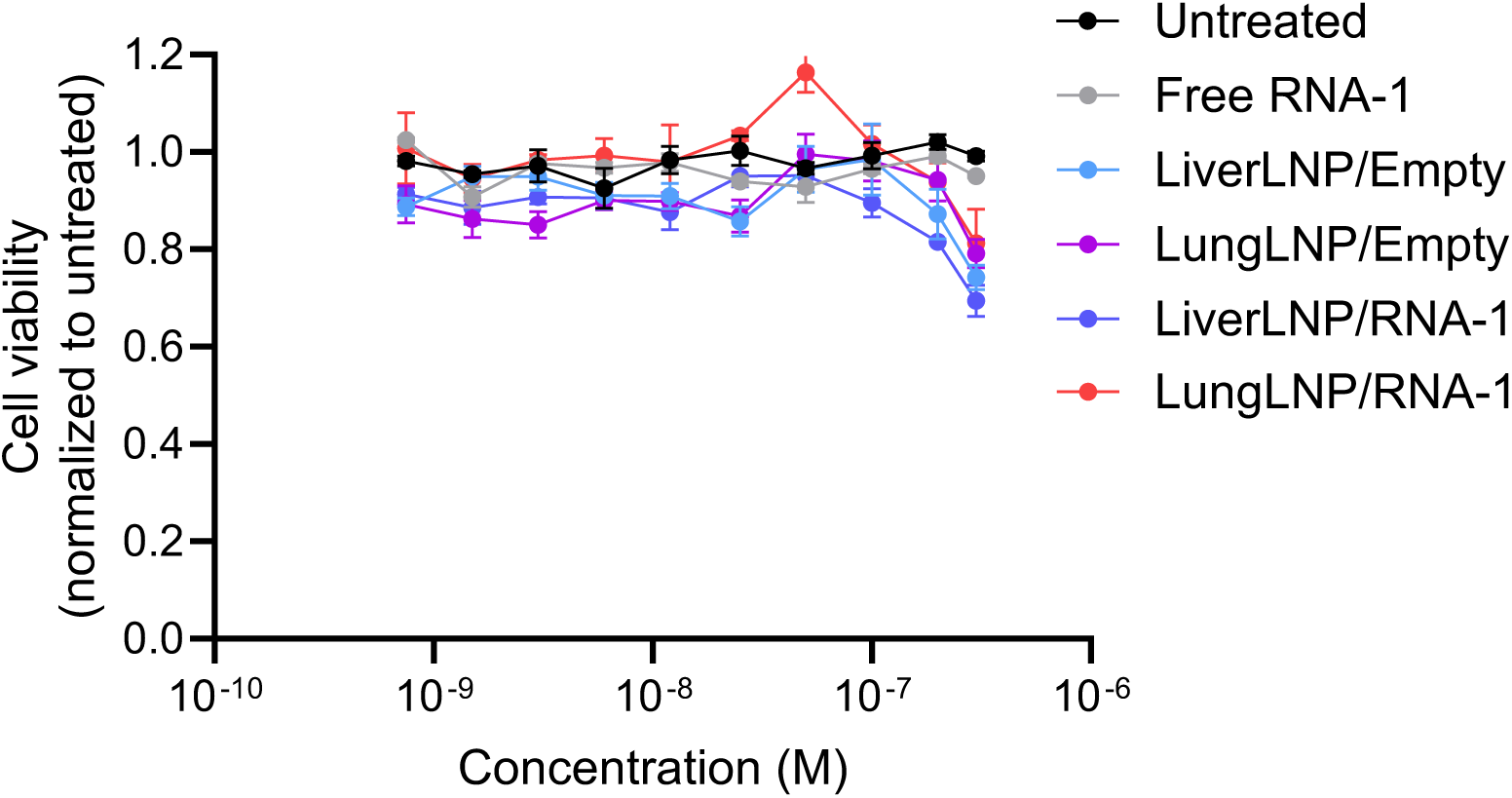
LNP/RNA-1 formulations maintain A549 cell viability. Cell viability of A549 cells treated with free RNA-1, LiverLNPs/RNA-1 or LungLNPs/RNA-1, or empty controls across indicated concentrations, normalized to untreated cells. Data presented as average ± SD, n = 3.

**Supplementary Fig. 3.**
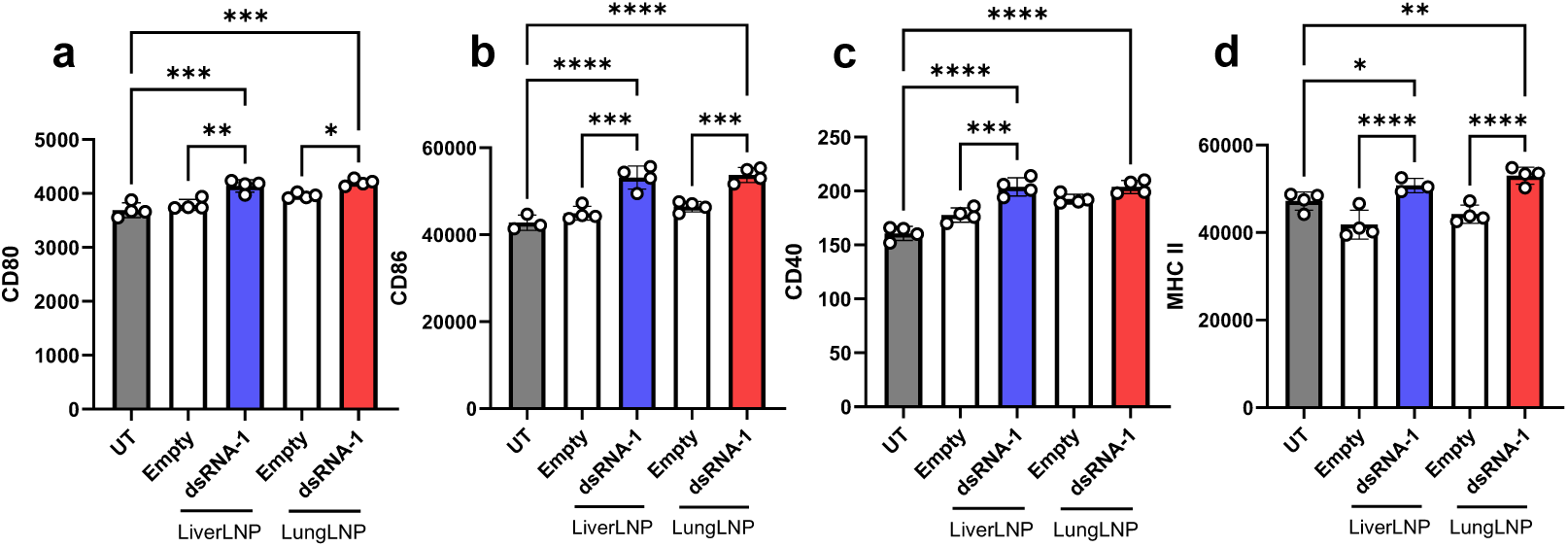
LNP/RNA-1 formulations activate bone marrow-derived dendritic cells in vitro. Bone marrow derived dendritic cells (BMDCs) were isolated from C57BL/6 mice and differentiated into DCs prior to treatment with free RNA-1, Liver-or LungLNPs/RNA-1, or empty LNP controls (RNA-1, 10nM). Surface expression of (a) CD80, (b) CD86, (c) CD40 and (d) MHC II was quantified by flow cytometry. LungLNP/RNA-1 and LiverLNP/RNA-1 have elicited comparable and marked upregulation of activation markers, whereas control formulations showed minimal effects. Data are represented as average ± SD, n=4.

**Supplementary Fig. 4.**
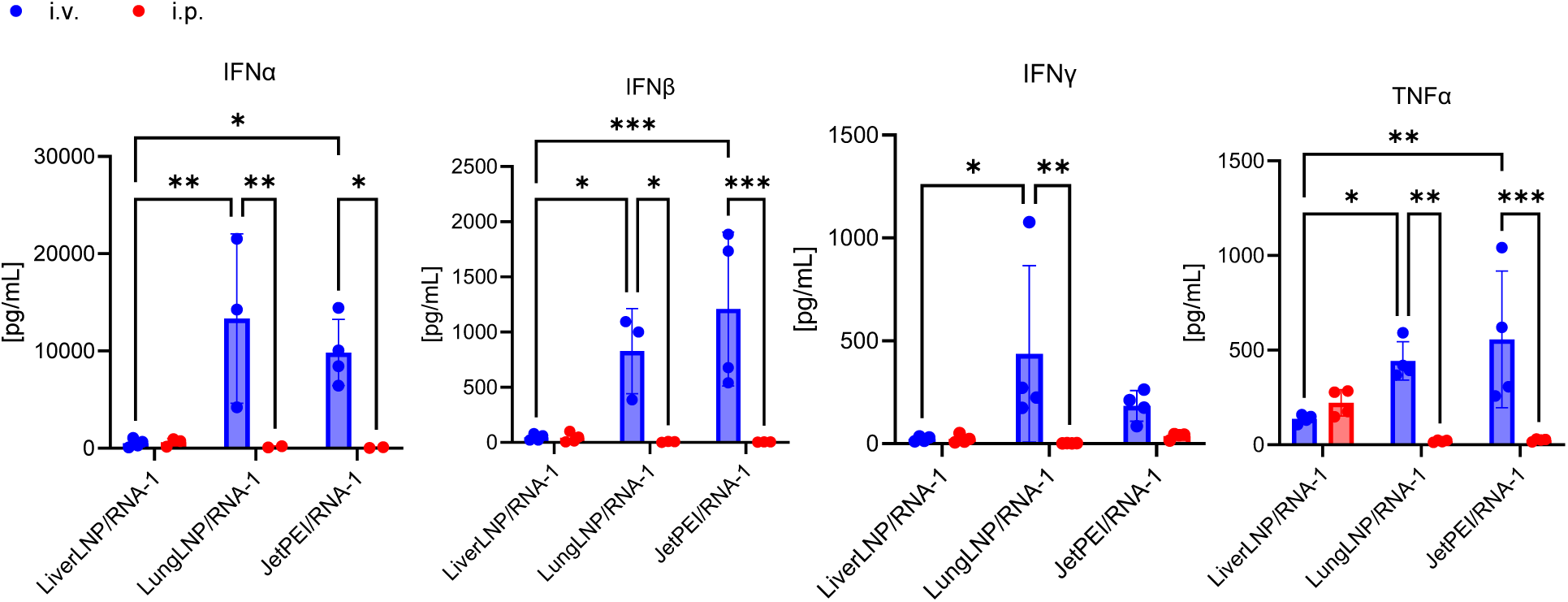
RNA-1 innate immune activation depends on formulation route of administration and BD. In vivo pharmacodynamic (PD) model measuring plasma cytokines following systemic (i.v.) or intraperitoneal (i.p.) administration of RNA-1 encapsulated in LiverLNPs, LungLNPs and JetPEI nanoparticles (2.2 mg/kg of RNA-1). Cytokine levels were measured at peak response (2h for IFNα, IFNβ, TNFα and 6h for IFNγ). Data are represented as mean ± SEM with n = 2–3. The data were analyzed by ordinary one-way ANOVA with Tukey’s multiple-comparisons test.

**Supplementary Fig. 5.**
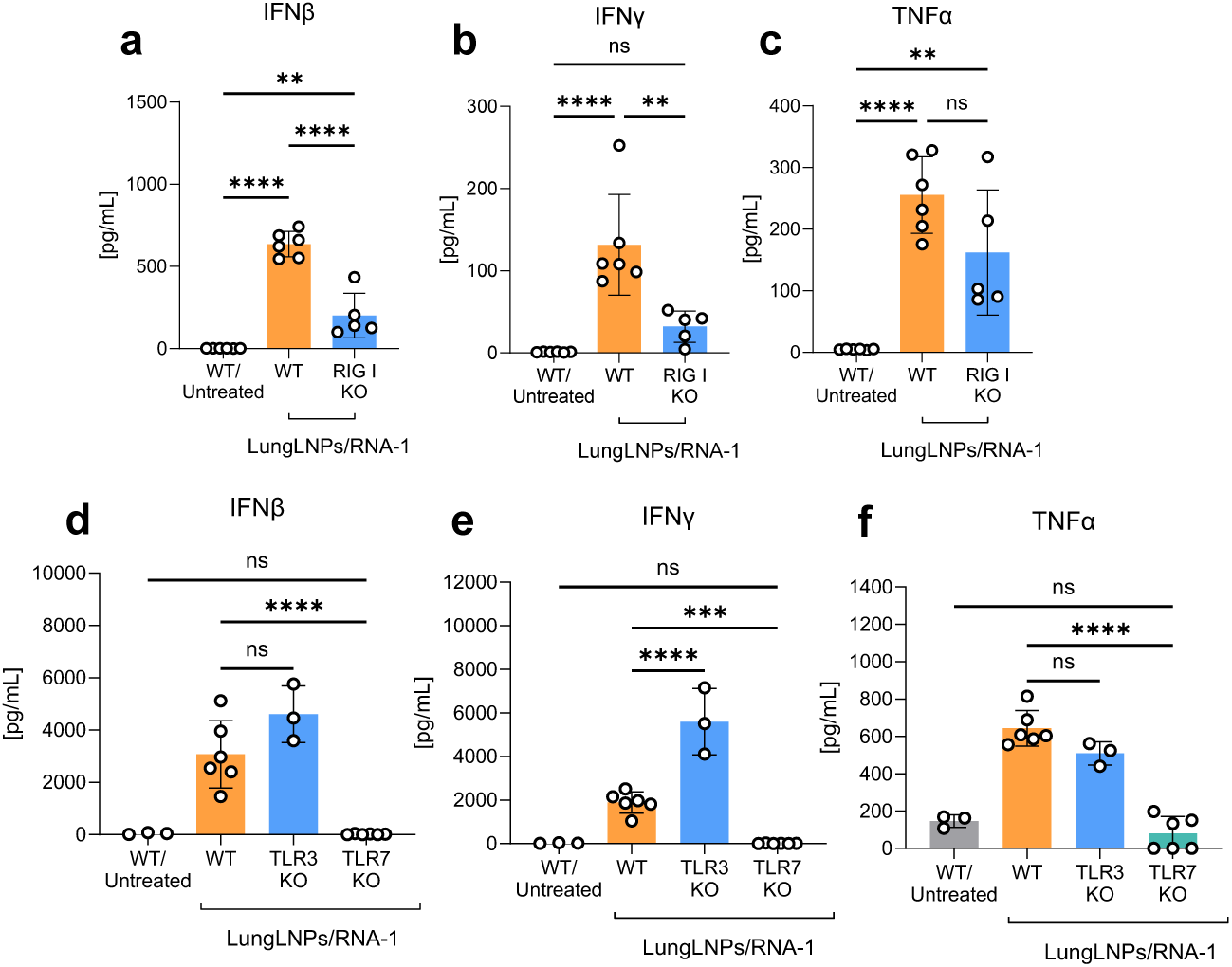
RNA-1-induced cytokine responses require TLR7 and RIG-I signaling. Knockout (KO) mice deficient in RNA-sensing pathways (TLR3, TLR7, or RIG-I) and wild-type (WT) controls were injected intravenously with LungLNP/RNA-1 (2.2 mg/kg RNA-1). Using the in vivo PD model study regimen, plasma cytokine levels were measured at 2 h, 6 h, and 24 h post-injection. Data shown represent plasma IFNβ, IFNγ, and TNFα induction at 2 h in (a–c) RIG-I KO mice; (d–f) TLR3 and TLR7 KO mice. Data are presented as mean ± SD with n = 3-6 biologically independent samples per group. Statistical analyses were performed using one-way ANOVA with Tukey’s multiple-comparisons test.

**Supplementary Fig. 6.**
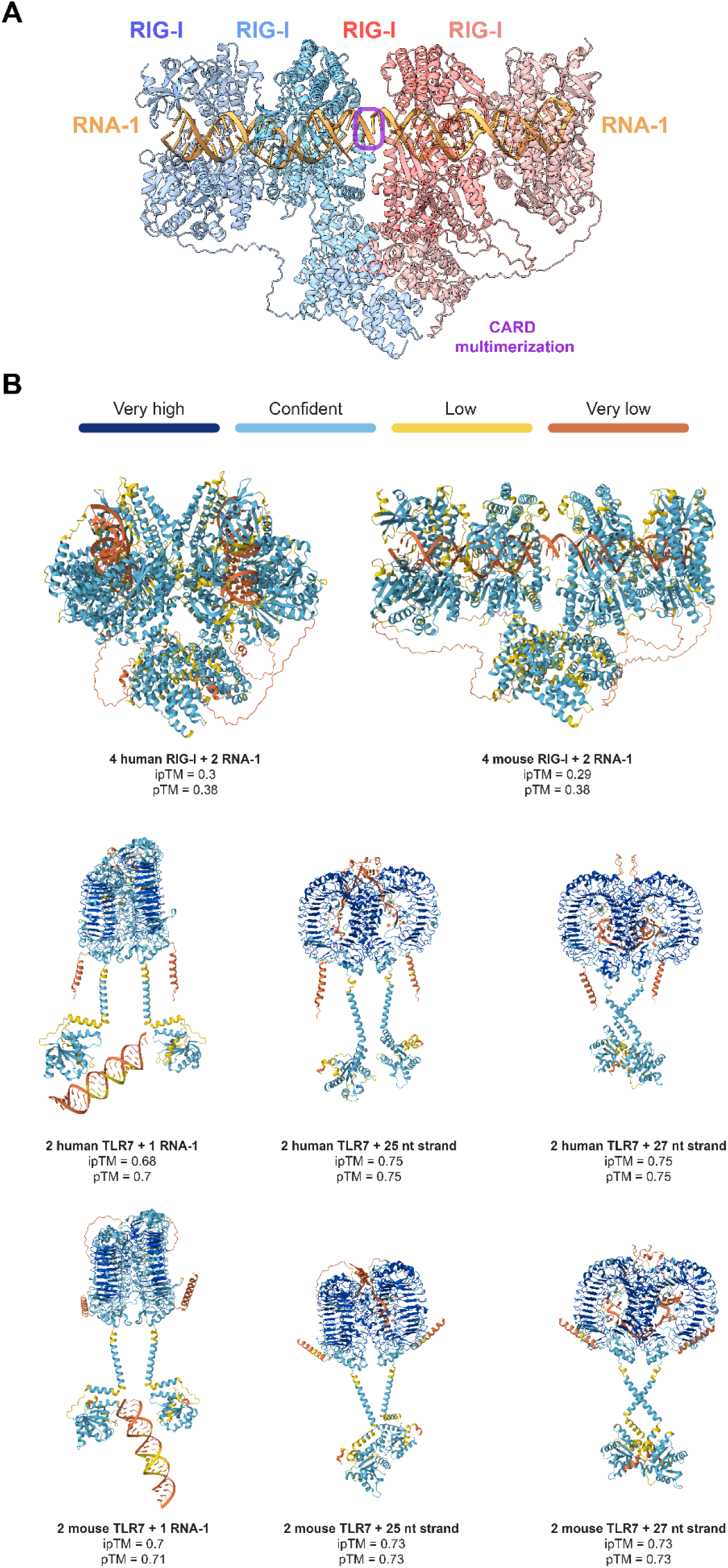
AlphaFold modeling reveals distinct RNA-1 binding modes for RIG-I and TLR7. (a) AlphaFold3 modeling of four mouse RIG-I with two copies of RNA-1, color-coded to indicate individual RIG-I proteins and RNA-1 duplexes. The putative GG-GG dimerization interaction between the two RNA-1 duplexes is outlined in purple. This AlphaFold prediction supports that two RNA-1 duplexes can accommodate the binding of four RIG-I and promote multimerization of their CARD domains in a lock-washer configuration, as required for downstream activation^1^. Prediction confidence levels for these models are shown in panel (b). (b) AlphaFold3 modeling of human and mouse RIG-I and TLR7 bound to either RNA-1 or its single-stranded components, color-coded by prediction confidence. Predicted template modeling (pTM) and interface predicted template modeling (ipTM) scores are shown for each prediction. Note that the large number of possible configurations of the RNA and protein interactions can result in lower confidence levels for any individual prediction. This modeling shows that while 4 RIG-I can bind to RNA-1 to multimerize, TLR7 dimers do not bind the duplex state but can bind the individual single strands.

**Supplementary Fig. 7.**
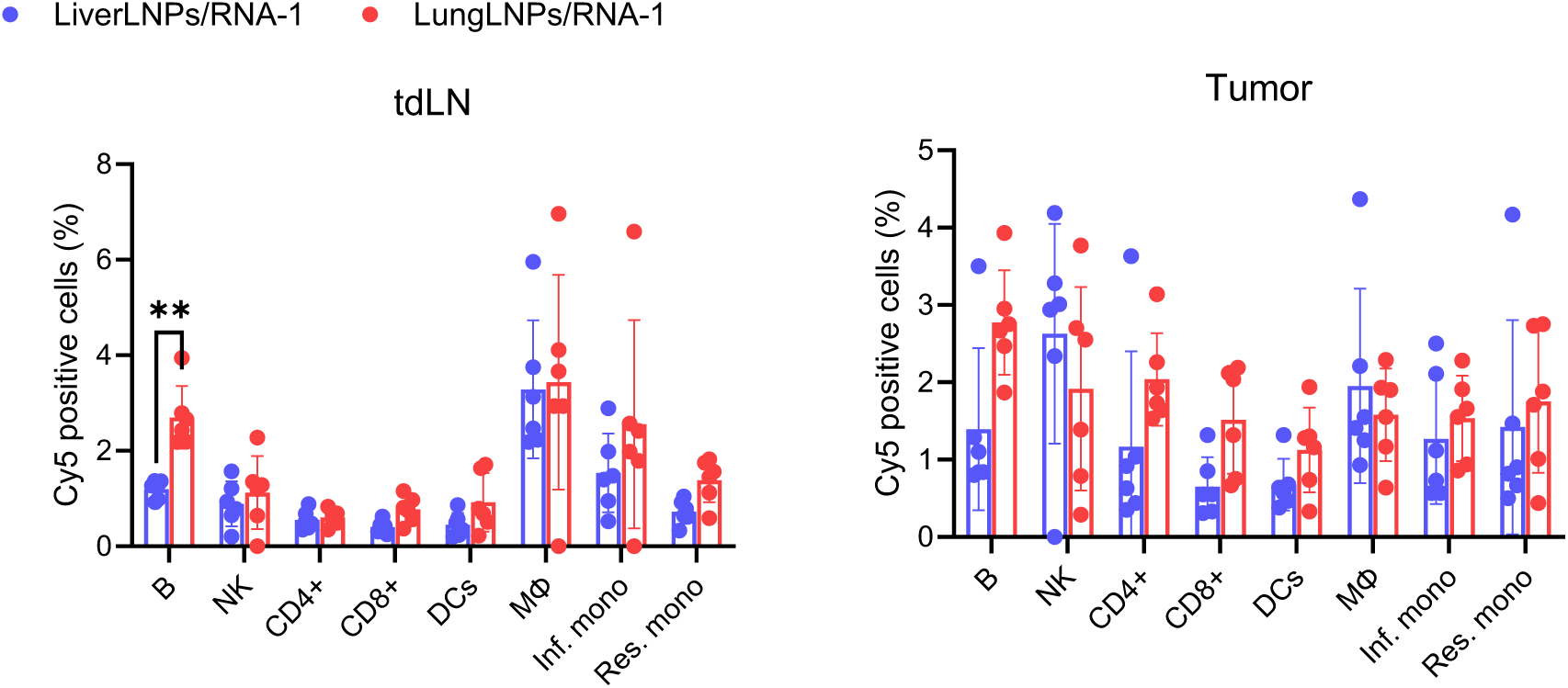
Flow cytometry analysis of cellular internalization of fluorescently labeled LungLNP/RNA-1^Cy5^ or LiverLNP/RNA-1^Cy5^ (2.2 mg/kg RNA-1) or PBS in mice bearing B16-F10 tumors LNPs in different tissues and immune cell populations in the tumor draining lymph node (tdLN) (A) and in the tumor (B).

**Supplementary Fig. 8.**
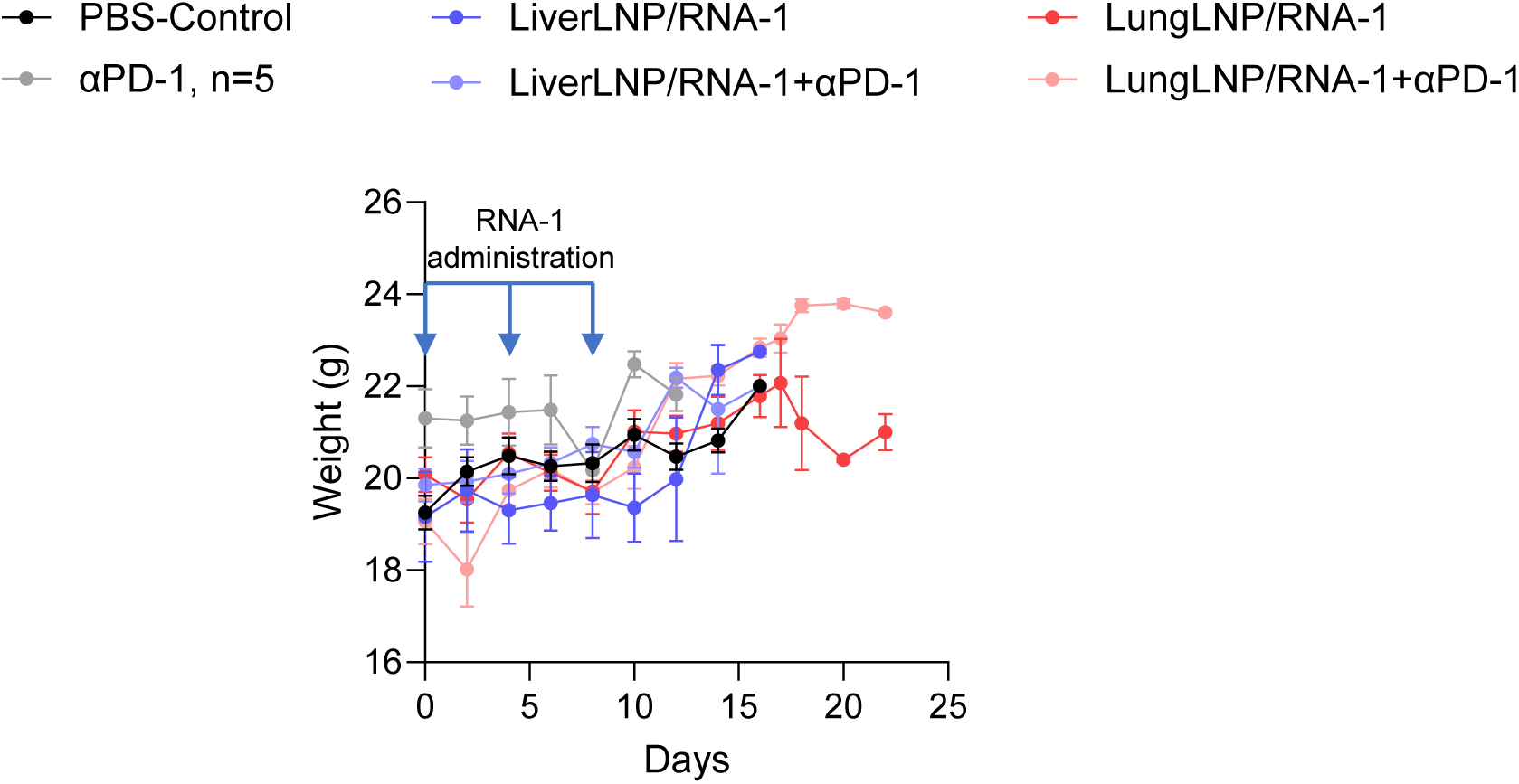
Mouse body weight tracking during a treatment regimen of LNP/RNA-1 therapy. Mice with B16F10 subcutaneous tumors were treated systemically with LungLNP/RNA-1 or LiverLNP /RNA-1 (2.2 mg/kg RNA-1) or control formulations, with or without intraperitoneal (i.p) aPD-1 antibody (100 μg), administered on days 7, 11, and 15 post tumor inoculation. The data represent mean ± s.e.m., from a representative experiment of two independent experiments with n = 5-10 biologically independent samples.

**Supplementary Fig. 9.**
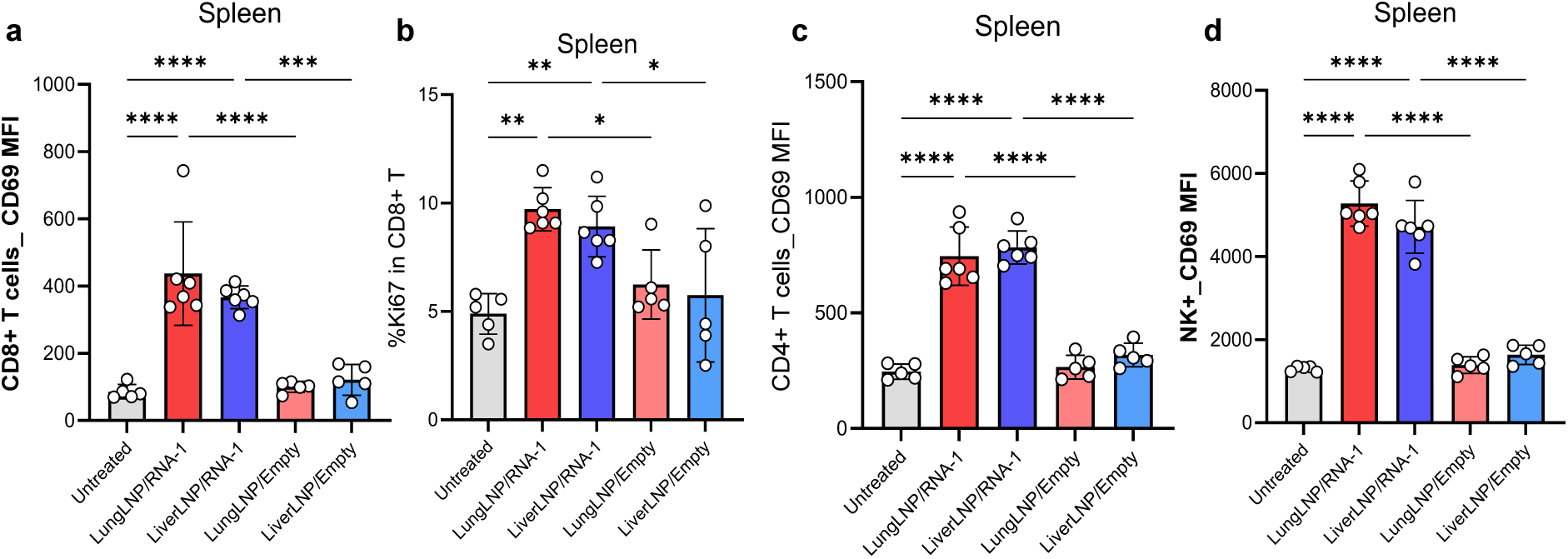
Both LungLNPs/RNA-1 and LiverLNPs/RNA-1 treatment activate lymphocytes within the spleen to the same extent. Flow cytometry analysis and quantification of (a) CD69 mean fluorescence intensity in CD8+ T cells (CD8+/NK1.1-/B220-/CD3+/CD11c-CD11b-/Ly6G-/CD45+), (b) precent of Ki67 positive CD8+ T cells, (c) CD69 mean fluorescence intensity in CD4+ T cells (CD4+/NK1.1-/B220-/CD3+/CD11c-CD11b-/Ly6G-/CD45+) and (d) CD69 mean fluorescence intensity in NK cells at the spleen 48h post systemic administration of RNA-1 using LungLNPs or LiverLNPs and corresponding empty nanoparticle controls and PBS control. The data represent mean ± SD with n = 5-6 biologically independent samples. The data were analyzed by one-way ANOVA with Tukey’s multiple comparisons test.

**Supplementary Fig. 10.**
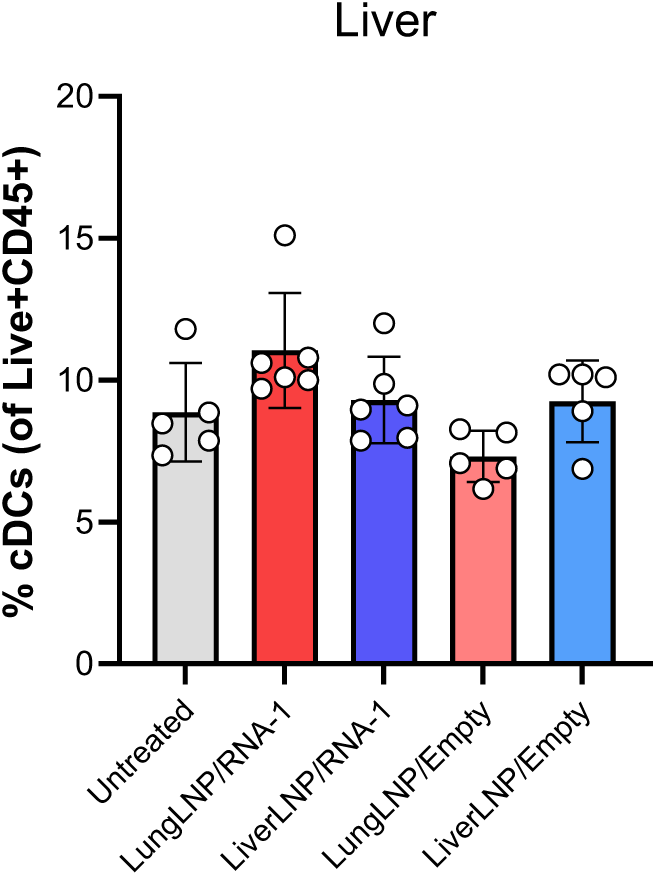
Infiltration of CD45+ immune cells wasn’t evident in the liver following treatment with RNA-1 LNPs. Flow cytometry analysis and quantification of % of CD45+ cells in the liver 48h post systemic administration of RNA-1 using LungLNPs or LiverLNPs and corresponding empty nanoparticle controls and PBS control. The data represent mean ± SD with n = 5-6 biologically independent samples. The data were analyzed by one-way ANOVA with Tukey’s multiple comparisons test.

**Supplementary Fig. 11.**
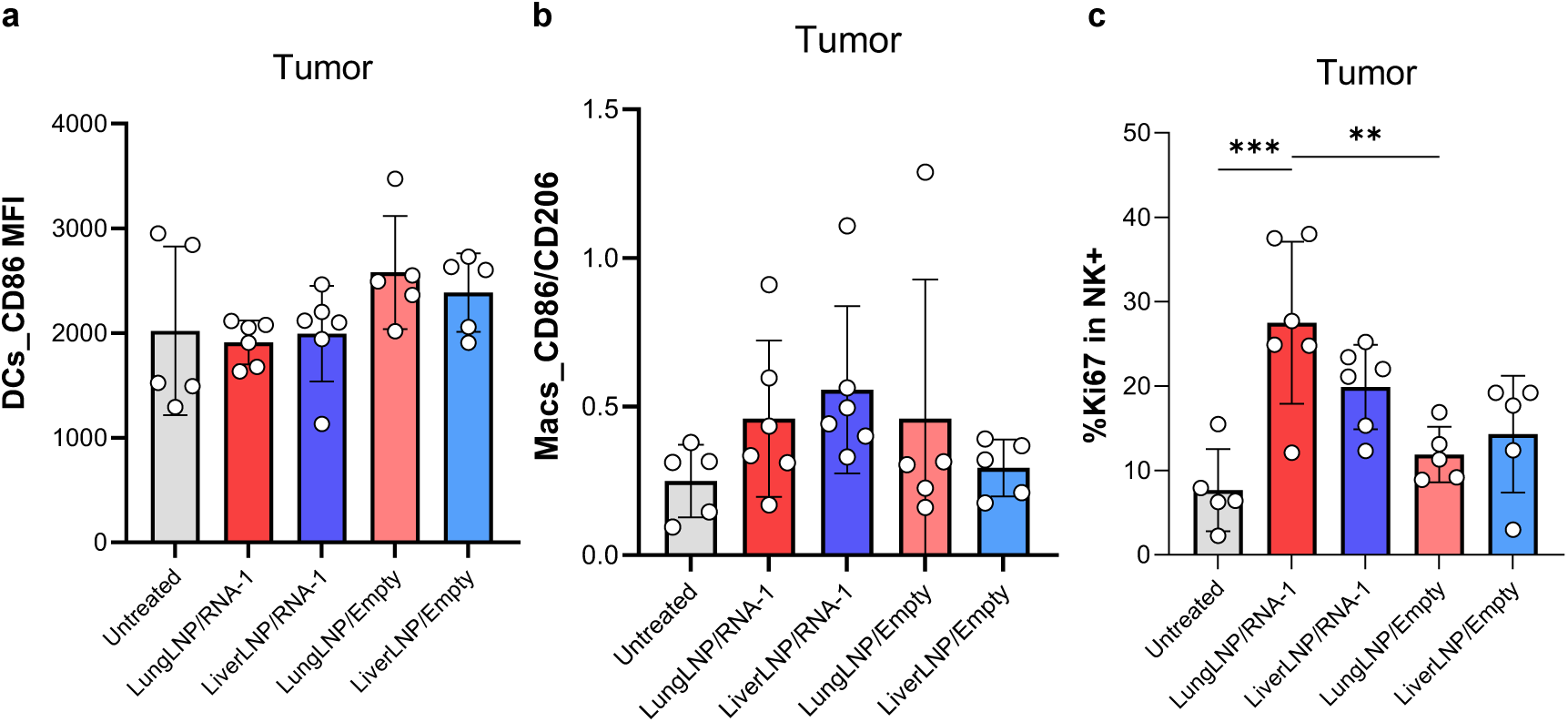
Dendritic cells (DCs) and macrophages (Macs) are not activated in the tumor while natural killer (NK) cells are specifically activated with LungLNPs/RNA-1. Flow cytometry analysis and quantification of CD86 mean fluorescence intensity in (a) DCs (F4/80-/CD11c+/Ly6G-/CD45+), (b) Macrophages (Macs) (F4/80+/CD11b+/Ly6G-/CD45+) and (c) precent of Ki67 positive NK cells (NK1.1+/B220-/CD3-/CD11c-CD11b-/Ly6G-/CD45+) at the TME 48h post systemic administration of RNA-1 using LungLNPs or LiverLNPs and corresponding empty nanoparticle controls and PBS control. The data represent mean ± SD with n = 5-6 biologically independent samples. The data were analyzed by one-way ANOVA with Tukey’s multiple comparisons test.

**Supplementary Fig. 12.**
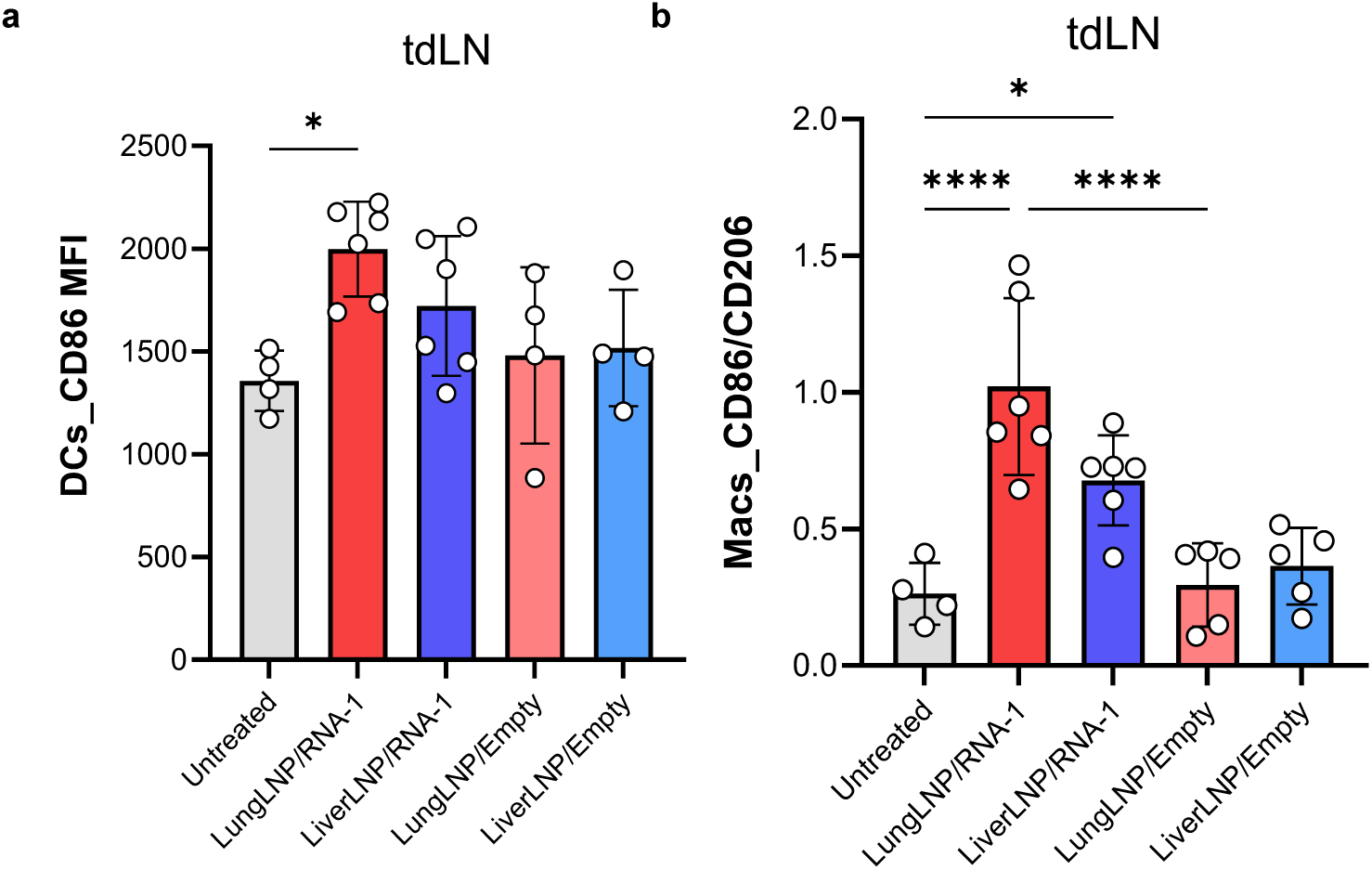
Dendritic cells (DCs) and macrophages (Macs) are activated in the tdLN specifically by LungLNPs/RNA-1 treatment. Flow cytometry analysis and quantification of (a) CD86 mean fluorescence intensity in DCs (F4/80-/CD11c+/Ly6G-/CD45+) and (b) CD86/CD206 mean fluorescence intensity in Macs (F4/80+/CD11b+/Ly6G-/CD45+) at the tdLN 48h post systemic administration of RNA-1 using LungLNPs or LiverLNPs and corresponding empty nanoparticle controls and PBS control. The data represent mean ± SD with n = 5-6 biologically independent samples. The data were analyzed by one-way ANOVA with Tukey’s multiple comparisons test.

**Supplementary Fig. 13:**
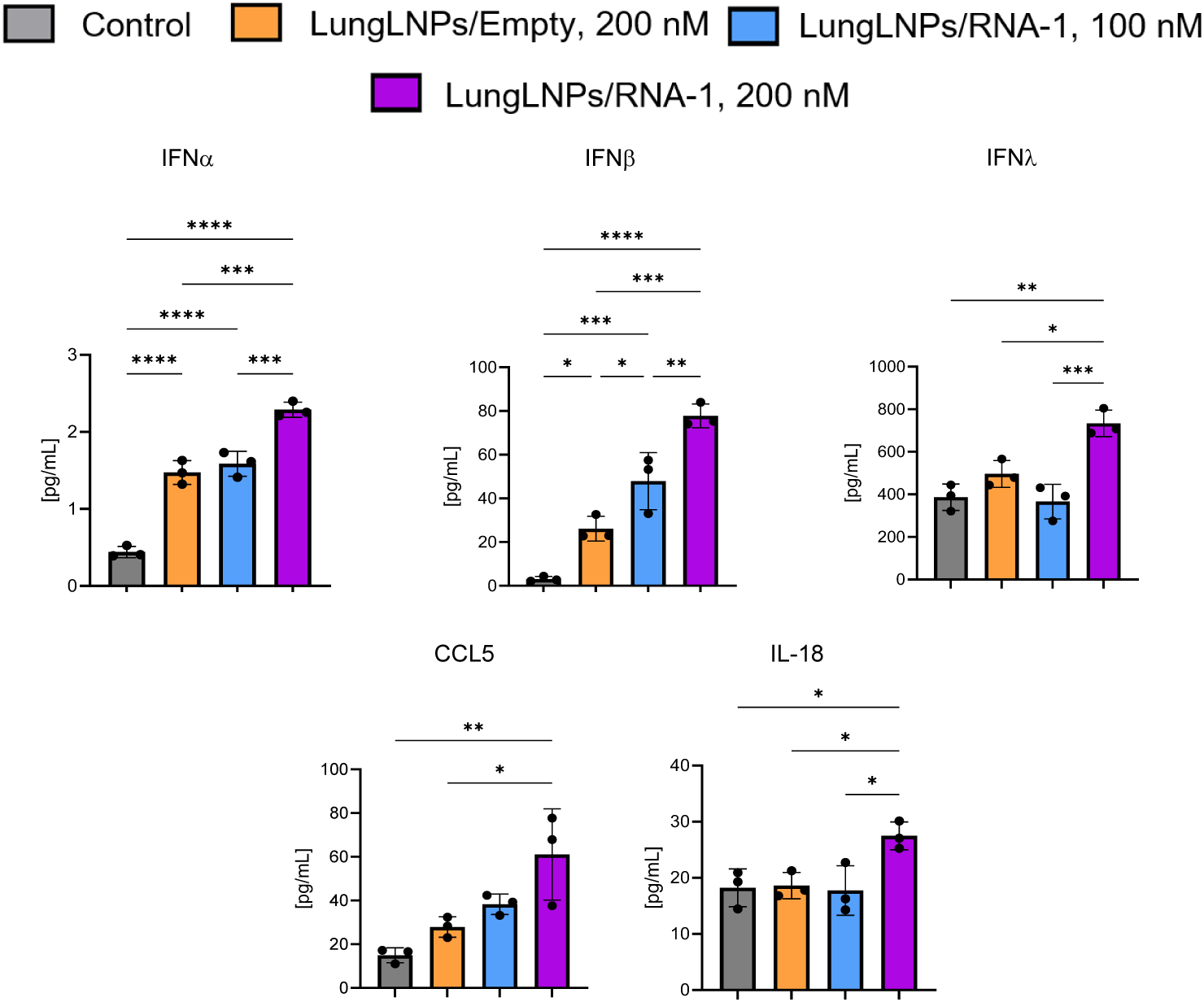
Human Lung cancer-chip studies reveals RNA-1 clinical translation potential in anticancer response. Quantification of cytokines, chemokines and ILs release at 2 h post first dosing according to the dosing regimen presented in Figure 6. The data were analyzed by one-way ANOVA with Tukey’s multiple-comparisons tests.

**Supplementary Fig 14.**
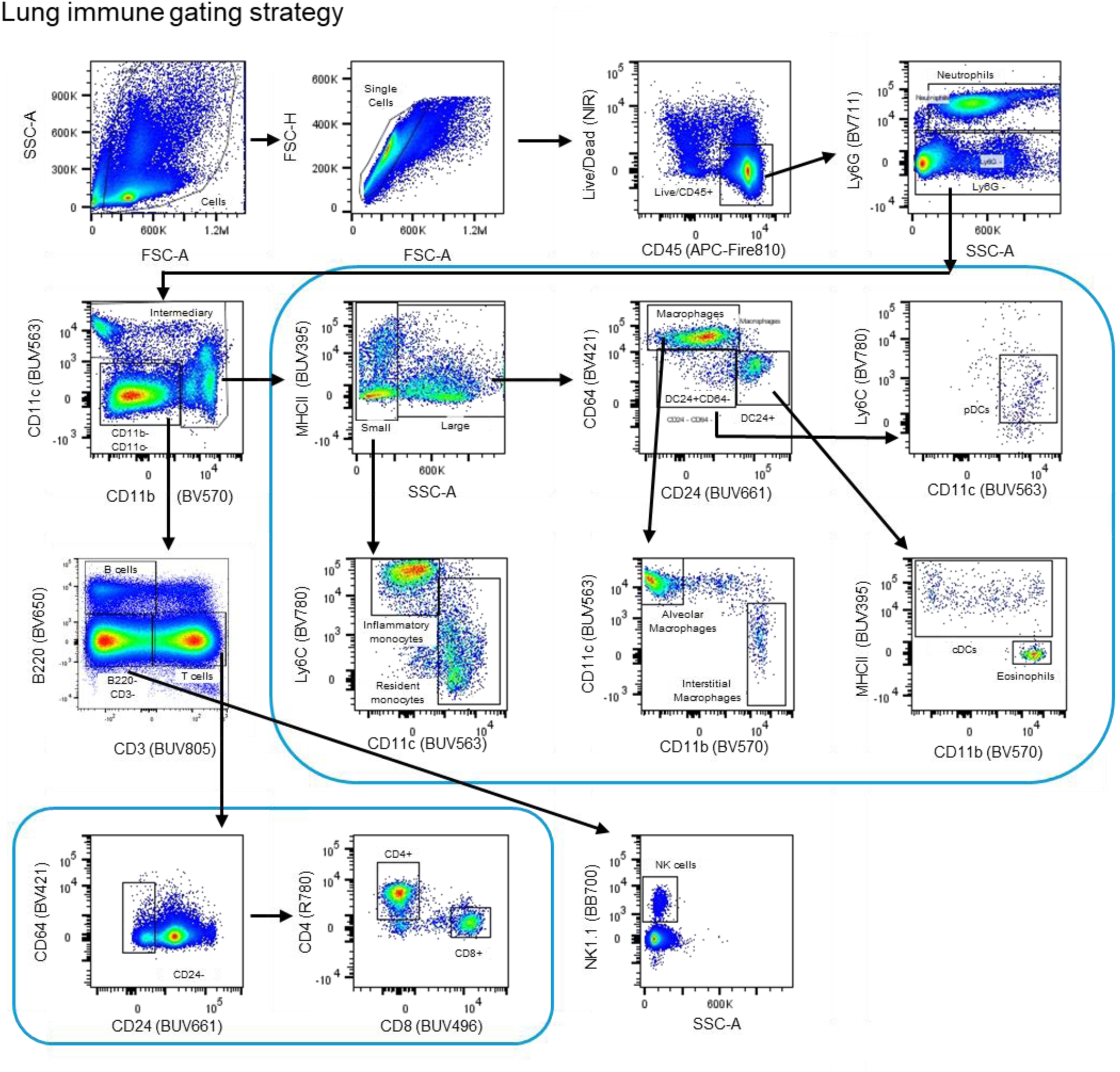
Gaiting strategy distinguishing immune cells populations in the lung based on fluorescent antibodies from Table S1.

**Supplementary Fig 15.**
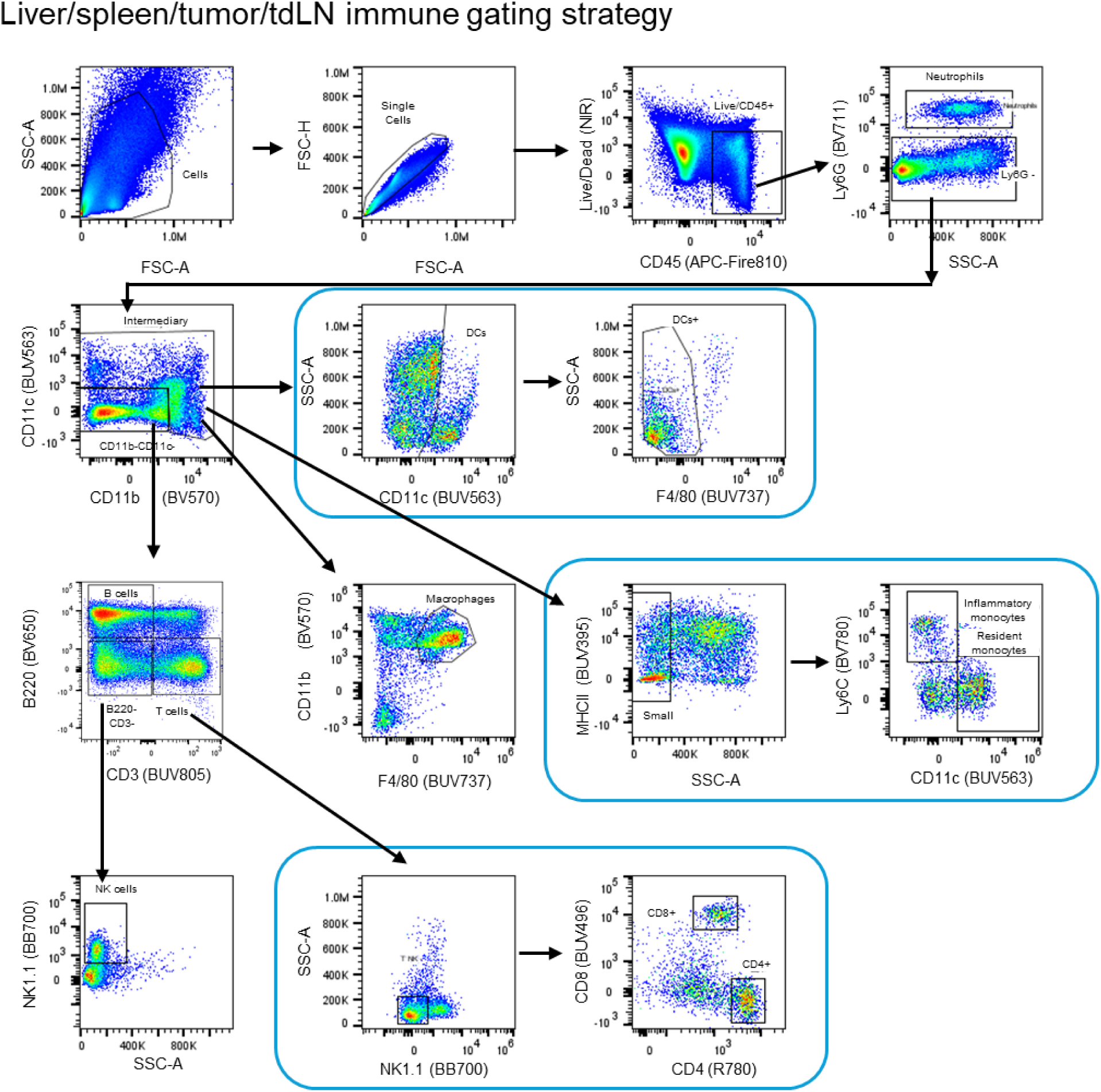
Gaiting strategy distinguishing immune cells populations in the liver/spleen/tumor/tdLN based on fluorescent antibodies from Table S1.

**Supplementary Fig 16.**
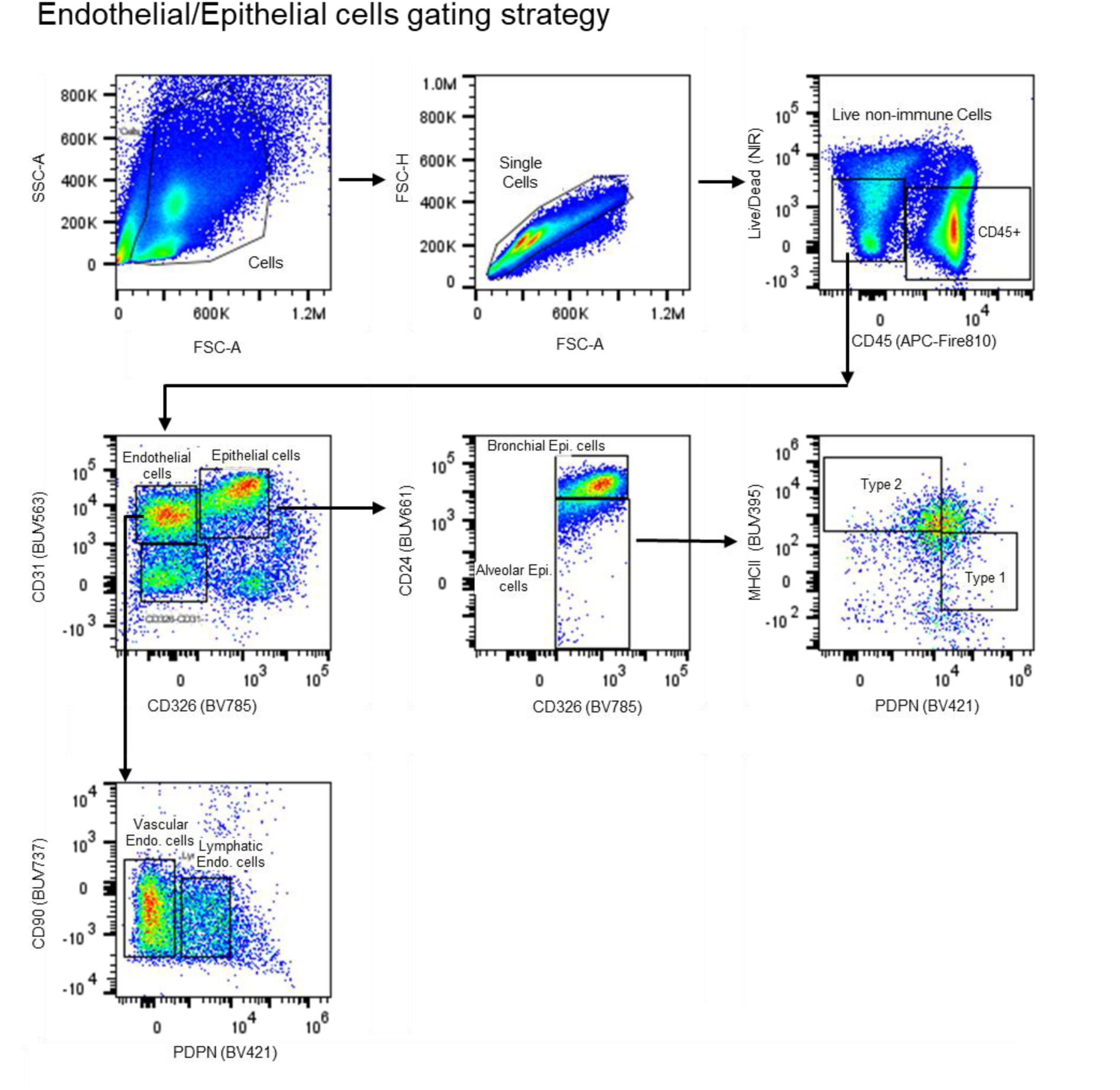
Gaiting strategy distinguishing endothelial and epithelial cells populations in the lung based on fluorescent antibodies from Table S1.

---

**Supplementary Table 1.** Markers used to define cell populations in mice organs using flow cytometry.

| **Marker** | **Fluorophore** | **Clone** | **Cat #** | **Supplier** |
| --- | --- | --- | --- | --- |
| LIVE/DEAD | LIVE/DEAD™ Fixable Near-IR Dead Cell Stain Kit, for 633 or 635 nm excitation | N/A | L10119 | Thermo Fisher |
| Ly6G | BV711 | 1A8 | 563979 | BD Biosciences |
| CD45 | APC/ Fire 810 | 30-F11 | 103173 | BioLegend |
| CD11b | BV570 | M1/70 | 101233 | BioLegend |
| CD11c | BUV563 | HL3 | 749091 | BD Biosciences |
| Ly6C | RB780 | HK1.4 rMAb | 755871 | BD Biosciences |
| MHCII | BUV395 | 2G9 | 569244 | BD Biosciences |
| B220 | BV650 | RA3-6B2 | 103241 | BioLegend |
| CD3 | BUV805 | 17A2 | 368-0032- 80 | Invitrogen |
| CD4 | R718 | GK1.5 | 567311 | BD Biosciences |
| CD8 | BUV496 | 53-6.7 | 364-0081- 82 | Invitrogen |
| NK1.1 | BB700 | PK136 | 566503 | BD Biosciences |
| CD64 | BV421 | X54-5/7.1 | 139309 | BioLegend |
| CD24 | BUV661 | 30-F11 | 752765 | BD Biosciences |
| f4/80 | BUV737 | T45-2342 (RUO) | 749283 | BD Biosciences |
| CD86 | BV785 | GL-1 | 105043 | BioLegend |
| CD206 | APC | C068C2 | 141708 | BioLegend |
| CD103 | BV605 | 2 E 7 | 121433 | BioLegend |
| KI67 | BV510 | B56 | 563462 | BD Biosciences |
| CD69 | PE/Dazzle 594 | H1.2F3 | 104536 | BioLegend |

## Supplementary Notes

**Supplementary note 1**: TLR7 and TLR8 are endosomal pattern-recognition receptors that are activated by single-stranded RNA species^2,3^. Although RNA-1 was administered as a self-dimerizing duplex, our data are consistent with literature findings that siRNAs such as RNA-1 may also be present as ssRNA that denatured in the endosome, where it could contribute to TLR7/8 activation^4,5^. At present, the origin of such ssRNA species in our system remains unclear. Possible explanations include partial duplex dissociation after uptake inside endosomes or the presence of minor ssRNA species before nanoparticle assembly. We note that cationic formulation-dependent destabilization of duplex RNA could play a role, such as in the case of DOTAP-containing formulations and JetPEI-based as suggested in the case of cationic ions and DNA destabilization^6^.

**Supplementary note 2**: Previous immunological studies^7,8^ demonstrate that endosomal TLR7 (and TLR9) signaling can directly induce the expression of cytosolic RNA sensors such as RIG-I in plasmacytoid dendritic cells (pDCs). In primary human pDCs, baseline RIG-I expression levels are low^9^, but stimulation with TLR7 agonists (e.g., imiquimod) or TLR9 ligands leads to rapid and dramatic upregulation of RIG-I, enabling these cells to subsequently sense cytosolic RNAs that otherwise cannot be detected under steady state. Crucially, this upregulation occurs in a type I IFN-independent manner, indicating that TLR engagement alone can reprogram pDCs to express RIG-I and integrate signals from both endosomal and cytosolic viral RNA sensing pathways^8^. This interplay correlates with the results in **Fig. 2 o-p** in mice KO models. In RIG-I knockout mice we demonstrate only a partial reduction in cytokine responses due to the contribution of the expressed TLR7 and their activation in these mice. On the other hand, in TLR7 KO mice, RNA-1 immune activation is completely abrogated although the expression of RIG I in these mice collectively is not affected. Thus, TLR7 might act as the dominant initiator of innate RNA sensing in this system, with RIG-I providing an additive or amplifying cytokines once its expression has been induced by TLR7 signaling. Dual agonism of TLR7 and RIG-I pathways as a strategy to enhance innate immune activation has precedent in the literature, where nanoparticle platforms co-deliver ligands targeting endosomal TLRs and RIG-I-like receptors to release type I IFN and pro-inflammatory cytokine responses^10^. In our case, RNA-1 itself appears to function as a dual agonist, engaging both TLR7-and RIG-I-dependent sensing within the same molecular structure.

## References

1. Vargason, A. M., Anselmo, A. C. & Mitragotri, S. The evolution of commercial drug delivery technologies. *Nat*. Biomed. Eng. 5, 951–967 (2021).

2. Hong, L., Li, W., Li, Y. & Yin, S. Nanoparticle-based drug delivery systems targeting cancer cell surfaces. RSC Adv. 13, 21365–21382 (2023).

3. Dosta, P. et al. Investigation of the enhanced antitumour potency of STING agonist after conjugation to polymer nanoparticles. Nat. Nanotechnol. 18, 1351–1363 (2023).

4. Dahis, D. et al. Focused Ultrasound Enhances Brain Delivery of Sorafenib Nanoparticles. Adv. NanoBiomed Res. 3, 2200142 (2023).

5. Luan, X., Wang, L., Song, G. & Zhou, W. Innate immune responses to RNA: sensing and signaling. Front. Immunol. 15, 1287940 (2024).

6. Marques, J. T., Meignin, C. & Imler, J.-L. An evolutionary perspective to innate antiviral immunity in animals. Cell Rep. 43, (2024).

7. Takeuchi, O. & Akira, S. Pattern recognition receptors and inflammation. Cell 140, 805–820 (2010).

8. Kumagai, Y. & Akira, S. Identification and functions of pattern-recognition receptors. J. Allergy Clin. Immunol. 125, 985–992 (2010).

9. Li, D. & Wu, M. Pattern recognition receptors in health and diseases. Signal Transduct. Target. Ther. 6, 291 (2021).

10. Jiang, Y. et al. Exploiting RIG-I-like receptor pathway for cancer immunotherapy. J. Hematol. Oncol.J Hematol Oncol 16, 8 (2023).

11. Stone, P. T. et al. Fabrication of RIG-I-Activating Nanoparticles for Intratumoral Immunotherapy via Flash Nanoprecipitation. Mol. Pharm. (2025).

12. Wang-Bishop, L. et al. Nanoparticle Retinoic Acid-Inducible Gene I Agonist for Cancer Immunotherapy. ACS Nano 18, 11631–11643 (2024).

13. Rodell, C. B. et al. TLR7/8-agonist-loaded nanoparticles promote the polarization of tumour-associated macrophages to enhance cancer immunotherapy. *Nat*. Biomed. Eng. 2, 578–588 (2018).

14. Kaur, A., Baldwin, J., Brar, D., Salunke, D. B. & Petrovsky, N. Toll-like receptor (TLR) agonists as a driving force behind next-generation vaccine adjuvants and cancer therapeutics. Curr. Opin. Chem. Biol. 70, 102172 (2022).

15. Chen, Y.-H., Wu, K.-H. & Wu, H.-P. Unraveling the complexities of toll-like receptors: from molecular mechanisms to clinical applications. Int. J. Mol. Sci. 25, 5037 (2024).

16. Huang, L., Ge, X., Liu, Y., Li, H. & Zhang, Z. The role of toll-like receptor agonists and their nanomedicines for tumor immunotherapy. Pharmaceutics 14, 1228 (2022).

17. Jiang, X. et al. Intratumoral delivery of RIG-I agonist SLR14 induces robust antitumor responses. J. Exp. Med. 216, 2854–2868 (2019).

18. Xu, H. et al. RIG-I RNA agonist activates immunostimulatory macrophages to enhance checkpoint immunotherapy for glioblastoma. BioRxiv Prepr. Serv. Biol. 2026.01.07.698153 (2026) doi:10.64898/2026.01.07.698153.

19. Rwandamuriye, F. X. et al. Local therapy with combination TLR agonists stimulates systemic anti-tumor immunity and sensitizes tumors to immune checkpoint blockade. Oncoimmunology 13, 2395067.

20. Mondal, J., Prabha, S., Griffith, T. S., Ferguson, D. & Panyam, J. Enhancing Cancer Therapy with TLR7/8 Agonists: Applications in Vaccines and Combination Treatments. Cancers 17, 3582 (2025).

21. Norris, P. A. & Kubes, P. Innate immunity of the lungs in homeostasis and disease. Mucosal Immunol. (2025).

22. Mettelman, R. C., Allen, E. K. & Thomas, P. G. Mucosal immune responses to infection and vaccination in the respiratory tract. Immunity 55, 749–780 (2022).

23. Diamond, M. S. & Kanneganti, T.-D. Innate immunity: the first line of defense against SARS-CoV-2. Nat. Immunol. 23, 165–176 (2022).

24. A tissue-scale strategy for sensing threats in barrier organs | bioRxiv. https://www.biorxiv.org/content/10.1101/2025.03.19.644134v1.abstract.

25. Si, L. et al. Self-assembling short immunostimulatory duplex RNAs with broad-spectrum antiviral activity. Mol. Ther. Nucleic Acids 29, 923–940 (2022).

26. Xu, J. et al. Identification of a natural viral RNA motif that optimizes sensing of viral RNA by RIG-I. MBio 6, 10.1128/mbio.01265-15 (2015).

27. Schlee, M. et al. Recognition of 5′ triphosphate by RIG-I helicase requires short blunt double-stranded RNA as contained in panhandle of negative-strand virus. Immunity 31, 25–34 (2009).

28. Iurescia, S., Fioretti, D. & Rinaldi, M. The innate immune signalling pathways: turning RIG-I sensor activation against cancer. Cancers 12, 3158 (2020).

29. Borden, E. C. Gene Regulation and Clinical Roles for Interferons in Neoplastic Diseases - Borden - 1998 - The Oncologist - Wiley Online Library. in.

30. Acebes-Fernández, V. et al. Nanomedicine and Onco-Immunotherapy: From the Bench to Bedside to Biomarkers. Nanomaterials 10, (2020).

31. Gisslinger, H. et al. Ropeginterferon alfa-2b versus standard therapy for polycythaemia vera (PROUD-PV and CONTINUATION-PV): a randomised, non-inferiority, phase 3 trial and its extension study. Lancet Haematol. 7, e196–e208 (2020).

32. Daunov, M. & Klisovic, R. B. Pegylated Interferons: Still a Major Player for the Treatment of Myeloproliferative Neoplasms. Am. Soc. Clin. Oncol. Educ. Book 45, e473912 (2025).

33. Healy, F. M., Dahal, L. N., Jones, J. R. E., Floisand, Y. & Woolley, J. F. Recent Progress in Interferon Therapy for Myeloid Malignancies. Front. Oncol. 11, (2021).

34. Krieg, A. M. New insights into the role of IFN-α/β and TLR7/8/9 in cancer immunotherapy and systemic autoimmunity. J. Immunother. Cancer 13, e012165 (2025).

35. Kirkwood, J. M. et al. Interferon Alfa-2b Adjuvant Therapy of High-Risk Resected Cutaneous Melanoma: The Eastern Cooperative Oncology Group Trial EST 1684. J. Clin. Oncol. 41, 425–435 (2023).

36. Marx, S. et al. RIG-I-induced innate antiviral immunity protects mice from lethal SARS-CoV-2 infection. Mol. Ther. Nucleic Acids 27, 1225–1234 (2022).

37. Tamir, H. et al. Induction of Innate Immune Response by TLR3 Agonist Protects Mice against SARS-CoV-2 Infection. Viruses 14, (2022).

38. Li, Y. et al. SARS-CoV-2 induces double-stranded RNA-mediated innate immune responses in respiratory epithelial-derived cells and cardiomyocytes. Proc. Natl. Acad. Sci. U. S. A. 118, e2022643118 (2021).

39. Cryer, A. M. et al. Restoration of cGAS in cancer cells promotes antitumor immunity via transfer of cancer cell–generated cGAMP. Proc. Natl. Acad. Sci. 122, e2409556122 (2025).

40. Momenzadeh, K. et al. Stimulation of fracture mineralization by salt-inducible kinase inhibitors. Front. Bioeng. Biotechnol. 12, 1450611 (2024).

41. Paranandi, K. S., Amar-Lewis, E., Mirkin, C. A. & Artzi, N. Nomadic Nanomedicines: Medicines Enabled by the Paracrine Transfer Effect. ACS Nano 19, 21–30 (2025).

42. Liu, S. et al. Charge-assisted stabilization of lipid nanoparticles enables inhaled mRNA delivery for mucosal vaccination. Nat. Commun. 15, 9471 (2024).

43. Peer, D. Induction of therapeutic gene silencing in leukocyte-implicated diseases by targeted and stabilized nanoparticles: A mini-review. J. Controlled Release 148, 63–68 (2010).

44. Tou, C. J. et al. Immune evasive DNA donors and recombinases license kilobase-scale writing. Nature 1–11 (2026) doi:10.1038/s41586-026-10241-z.

45. Akinc, A. et al. A combinatorial library of lipid-like materials for delivery of RNAi therapeutics. Nat. Biotechnol. 26, 561–569 (2008).

46. Dilliard, S. A., Cheng, Q. & Siegwart, D. J. On the mechanism of tissue-specific mRNA delivery by selective organ targeting nanoparticles. Proc. Natl. Acad. Sci. 118, e2109256118 (2021).

47. Moore, S. T. et al. Multiplexed lipid nanoparticle barcoding reveals tissue-dynamic kinetic insights and enriched cellular tropism in hepatic zones. Nat. Commun. (2026).

48. Amar-Lewis, E. et al. Quaternized starch-based carrier for siRNA delivery: From cellular uptake to gene silencing. J. Controlled Release 185, 109–120 (2014).

49. Cheng, Q. et al. Selective organ targeting (SORT) nanoparticles for tissue-specific mRNA delivery and CRISPR–Cas gene editing. Nat. Nanotechnol. 15, 313–320 (2020).

50. Wang, X. et al. Preparation of selective organ-targeting (SORT) lipid nanoparticles (LNPs) using multiple technical methods for tissue-specific mRNA delivery. Nat. Protoc. 18, 265–291 (2023).

51. McNab, F., Mayer-Barber, K., Sher, A., Wack, A. & O’garra, A. Type I interferons in infectious disease. Nat. Rev. Immunol. 15, 87–103 (2015).

52. Kawai, T. & Akira, S. The role of pattern-recognition receptors in innate immunity: update on Toll-like receptors. Nat. Immunol. 11, 373–384 (2010).

53. Poster: in vivo-jetPEI®. https://www.polyplus-sartorius.com/poster-in-vivo-jetpei-an-alternative-to-lipid-based-reagents-or-viral-vectors-for-nucleic-acid-mediated-therapies (2023).

54. Amar-Lewis, E. et al. Quaternized Starch-Based Composite Nanoparticles for siRNA Delivery to Tumors. ACS Appl. Nano Mater. 4, 2218–2229 (2021).

55. Amar-Lewis, E. et al. Elucidating siRNA Cellular Delivery Mechanism Mediated by Quaternized Starch Nanoparticles. Small 20, 2405524 (2024).

56. Goula, D. et al. Polyethylenimine-based intravenous delivery of transgenes to mouse lung. Gene Ther. 5, 1291–1295 (1998).

57. Kwak, G., Lee, D. & Suk, J. S. Advanced approaches to overcome biological barriers in respiratory and systemic routes of administration for enhanced nucleic acid delivery to the lung. Expert Opin. Drug Deliv. 20, 1531–1552 (2023).

58. Marques, J. T. & Williams, B. R. G. Activation of the mammalian immune system by siRNAs. Nat. Biotechnol. 23, 1399–1405 (2005).

59. Kaushal, A. Innate immune regulations and various siRNA modalities. Drug Deliv. Transl. Res. 13, 2704–2718 (2023).

60. Ren, Y. et al. Impact of ionizable lipid type on the pharmacokinetics and biodistribution of mRNA-lipid nanoparticles after intravenous and subcutaneous injection. J. Control. Release Off. J. Control. Release Soc. 384, 113945 (2025).

61. Hosseini-Kharat, M., Bremmell, K. E. & Prestidge, C. A. Why do lipid nanoparticles target the liver? Understanding of biodistribution and liver-specific tropism. Mol. Ther. Methods Clin. Dev. 33, (2025).

62. Liu, Y. et al. Development of mRNA Lipid Nanoparticles: Targeting and Therapeutic Aspects. Int. J. Mol. Sci. 25, (2024).

63. Kon, E., Ad-El, N., Hazan-Halevy, I., Stotsky-Oterin, L. & Peer, D. Targeting cancer with mRNA–lipid nanoparticles: key considerations and future prospects. Nat. Rev. Clin. Oncol. 20, 739–754 (2023).

64. Ramos-Gonzalez, M. R., Vazquez-Garza, E., Garcia-Rivas, G., Rodriguez-Aguayo, C. & Chavez-Reyes, A. Therapeutic Effects of WT1 Silencing via Respiratory Administration of Neutral DOPC Liposomal-siRNA in a Lung Metastasis Melanoma Murine Model. Non-Coding RNA 9, (2023).

65. Freeman, F. E. et al. Localized nanoparticle-mediated delivery of miR-29b normalizes the dysregulation of bone homeostasis caused by osteosarcoma whilst simultaneously inhibiting tumor growth. Adv. Mater. 35, 2207877 (2023).

66. Fekete, T. et al. Human plasmacytoid and monocyte-derived dendritic cells display distinct metabolic profile upon RIG-I activation. Front. Immunol. 9, 3070 (2018).

67. Toy, R. et al. TLR7 and RIG-I dual-adjuvant loaded nanoparticles drive broadened and synergistic responses in dendritic cells in vitro and generate unique cellular immune responses in influenza vaccination. J. Controlled Release 330, 866–877 (2021).

68. Bruni, D. et al. Viral entry route determines how human plasmacytoid dendritic cells produce type I interferons. Sci. Signal. 8, ra25 (2015).

69. Hassell, B. A. et al. Human Organ Chip Models Recapitulate Orthotopic Lung Cancer Growth, Therapeutic Responses, and Tumor Dormancy In Vitro. Cell Rep. 21, 508–516 (2017).

70. Dasgupta, Q. et al. A human lung alveolus-on-a-chip model of acute radiation-induced lung injury. Nat. Commun. 14, 6506 (2023).

71. Plebani, R. et al. Modeling pulmonary cystic fibrosis in a human lung airway-on-a-chip. J. Cyst. Fibros. 21, 606–615 (2022).

72. RIGImmune Inc. A Two-Part Randomized, Double-Blind Placebo Controlled Trial to Assess the Safety and Tolerability of Single and Repeat Ascending Intranasal Doses of RIG-101 in Healthy Participants Followed by Repeat Daily Administration in Adult Participants With Asthma [Part A] Followed by a Randomized Double-Blind Placebo Controlled Part to Assess the Efficacy and Safety of RIG-101 in Adult Participants With Asthma Before and After Viral Challenge With Human Rhinovirus RV-A16 [Part B]. https://clinicaltrials.gov/study/NCT07488897 (2026).

73. Merck Sharp &Dohme LLC. A Phase 1&#x2F;1b, Open-Label Clinical Study of Intratumoral&#x2F;Intralesional Administration of MK-4621&#x2F;JetPEI as Monotherapy or in Combination With Pembrolizumab (MK-3475) in Participants With Advanced&#x2F;Metastatic or Recurrent Solid Tumors. https://clinicaltrials.gov/study/NCT03739138 (2022).

74. Merck Sharp &Dohme LLC. A Phase I&#x2F;II, Multicenter, Open-Label, Clinical Trial of Intratumoral&#x2F;Intralesional Administration of RGT100 in Subjects With Advanced or Recurrent Tumors. https://clinicaltrials.gov/study/NCT03065023 (2019).

75. Van Lysebetten, D. et al. Lipid-Polyglutamate Nanoparticle Vaccine Platform. ACS Appl. Mater. Interfaces 13, 6011–6022 (2021).

76. Battistella, C. et al. Delivery of Immunotherapeutic Nanoparticles to Tumors via Enzyme-Directed Assembly. Adv. Healthc. Mater. 8, e1901105 (2019).

77. Saitoh, S.-I. et al. TLR7 mediated viral recognition results in focal type I interferon secretion by dendritic cells. Nat. Commun. 8, 1592 (2017).

78. McKenna, K., Beignon, A.-S. & Bhardwaj, N. Plasmacytoid dendritic cells: linking innate and adaptive immunity. J. Virol. 79, 17–27 (2005).

79. Kohli, K., Pillarisetty, V. G. & Kim, T. S. Key chemokines direct migration of immune cells in solid tumors. Cancer Gene Ther. 29, 10–21 (2022).

80. Busselaar, J., Sijbranda, M. & Borst, J. The importance of type I interferon in orchestrating the cytotoxic T-cell response to cancer. Immunol. Lett. 270, 106938 (2024).

81. Elia, U. & Peer, D. A symphony of immunity: How HPV mRNA-LNP vaccination orchestrates systemic anti-tumor responses. Mol. Ther. Nucleic Acids 37, 102823 (2026).

82. Dosta, P., Cryer, A. M., Prado, M. & Artzi, N. Bioengineering strategies to optimize STING agonist therapy. Nat. Rev. Bioeng. 3, 660–680 (2025).

## References

1. Artzi, N., et al. Lipid nanoparticles for the treatment of vascular diseases. (2025).

2. Cryer, A. M. et al. Restoration of cGAS in cancer cells promotes antitumor immunity via transfer of cancer cell–generated cGAMP. Proc. Natl. Acad. Sci. 122, e2409556122 (2025).

3. Dosta, P. et al. Delivery of Stimulator of Interferon Genes (STING) Agonist Using Polypeptide-Modified Dendrimer Nanoparticles in the Treatment of Melanoma. Adv. NanoBiomed Res. 1, 2100006 (2021).

4. Wang-Bishop, L. et al. Nanoparticle Retinoic Acid-Inducible Gene I Agonist for Cancer Immunotherapy. ACS Nano 18, 11631–11643 (2024).

5. Baumann, Z. et al. Optimized full-spectrum flow cytometry panel for deep immunophenotyping of murine lungs. *Cell Rep*. Methods 4, (2024).

6. Hassell, B. A. et al. Human Organ Chip Models Recapitulate Orthotopic Lung Cancer Growth, Therapeutic Responses, and Tumor Dormancy In Vitro. Cell Rep. 21, 508–516 (2017).

## References

1. Peisley, A., Wu, B., Xu, H., Chen, Z. J. & Hur, S. Structural basis for ubiquitin-mediated antiviral signal activation by RIG-I. Nature 509, 110–114 (2014).

2. Kaushal, A. Innate immune regulations and various siRNA modalities. Drug Deliv. Transl. Res. 13, 2704–2718 (2023).

3. Sioud, M. Single-stranded small interfering RNA are more immunostimulatory than their double-stranded counterparts: A central role for 2′-hydroxyl uridines in immune responses. Eur. J. Immunol. 36, 1222–1230 (2006).

4. Marques, J. T. & Williams, B. R. G. Activation of the mammalian immune system by siRNAs. Nat. Biotechnol. 23, 1399–1405 (2005).

5. Goodchild, A. et al. Sequence determinants of innate immune activation by short interfering RNAs. BMC Immunol. 10, 40 (2009).

6. Zhang, C. et al. Counterintuitive DNA destabilization by monovalent salt at high concentrations due to overcharging. Nat. Commun. 16, 113 (2025).

7. Szabo, A. et al. TLR ligands upregulate RIG-I expression in human plasmacytoid dendritic cells in a type I IFN-independent manner. Immunol. Cell Biol. 92, 671–678 (2014).

8. Fekete, T. et al. Human Plasmacytoid and Monocyte-Derived Dendritic Cells Display Distinct Metabolic Profile Upon RIG-I Activation. Front. Immunol. 9, (2018).

9. Bruni, D. et al. Viral entry route determines how human plasmacytoid dendritic cells produce type I interferons. Sci. Signal. 8, ra25–ra25 (2015).

10. Toy, R. et al. TLR7 and RIG-I dual-adjuvant loaded nanoparticles drive broadened and synergistic responses in dendritic cells *in vitro* and generate unique cellular immune responses in influenza vaccination. J. Controlled Release 330, 866–877 (2021).

